# Perinatal High Fat Diet Exposure Alters Oxytocin and Corticotropin Releasing Factor Inputs onto Vagal Neurocircuits Controlling Gastric Motility

**DOI:** 10.1101/2022.11.28.517564

**Authors:** Kaitlin E. Carson, Jared Alvarez, Jasmine Mackley, R. Alberto Travagli, Kirsteen N. Browning

## Abstract

Perinatal high fat diet (pHFD) exposure alters the development of vagal neurocircuits that control gastrointestinal (GI) motility and reduce stress resiliency in offspring. Descending oxytocin (OXT; prototypical anti-stress peptide) and corticotropin releasing factor (CRF; prototypical stress peptide) inputs from the paraventricular nucleus (PVN) of the hypothalamus to the dorsal motor nucleus of the vagus (DMV) modulate the GI stress response. How these descending inputs, and their associated changes to GI motility and stress responses, are altered following pHFD exposure are, however, unknown. The present study used retrograde neuronal tracing experiments, *in vivo* recordings of gastric tone, motility, and gastric emptying rates, and *in vitro* electrophysiological recordings from brainstem slice preparations to investigate the hypothesis that pHFD alters descending PVN-DMV inputs and dysregulates vagal brain-gut responses to stress. Compared to controls, rats exposed to pHFD had slower gastric emptying rates and did not respond to acute stress with the expected delay in gastric emptying. Neuronal tracing experiments demonstrated that pHFD reduced the number of PVN^OXT^ neurons that project to the DMV, but increased PVN^CRF^ neurons. Both *in vitro* electrophysiology recordings of DMV neurons and *in vivo* recordings of gastric motility and tone demonstrated that, following pHFD, PVN^CRF^-DMV projections were tonically active, and that pharmacological antagonism of brainstem CRF1 receptors restored the appropriate gastric response to brainstem OXT application. These results suggest that pHFD exposure disrupts descending PVN-DMV inputs, leading to a dysregulated vagal brain-gut response to stress.

**Summary Figure:** 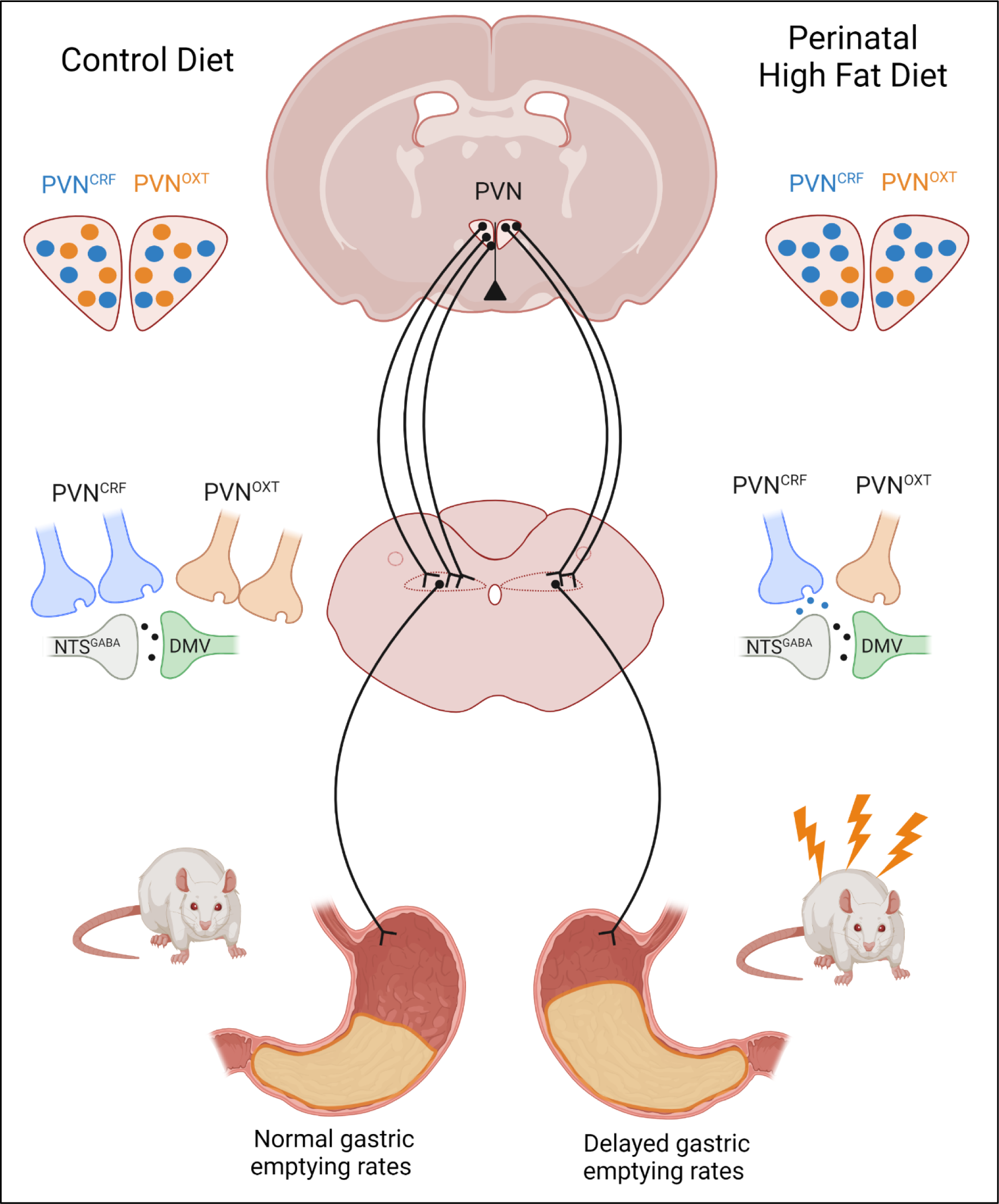

**Keypoints:**

- Maternal high fat diet exposure is associated with gastric dysregulation and stress sensitivity in offspring
- The present study demonstrates that perinatal high fat diet exposure downregulates hypothalamic-vagal oxytocin (OXT) inputs but upregulates hypothalamic-vagal corticotropin releasing factor (CRF) inputs
- Both in vitro and in vivo studies demonstrated that, following perinatal high fat diet, CRF receptors were tonically active at NTS-DMV synapses, and that pharmacological antagonism of these receptors restored appropriate gastric response to OXT
- The current study suggests that perinatal high fat diet exposure disrupts descending PVN-DMV inputs, leading to a dysregulated vagal brain-gut response to stress.

## Introduction

Obesity is a significant and growing problem that has resulted in an increase in maternal obesity, with approximately 30% of mothers in the United States in 2019 being classified as obese (Driscoll, 2020). Although multifactorial in origin, involving both genetic and environmental factors (Levin, 2006, 2010), obesity ultimately results from an imbalance of caloric intake relative to expenditure, and the ready availability of calorically dense, high fat foods has significantly worsened this problem (Bray *et al*., 2004).

Consumption of a high fat diet (HFD) has well known, major consequences to metabolic health, however there is also evidence that HFD exposure *in utero* and during early postnatal life (i.e., the perinatal period) can profoundly affect offspring neural and metabolic development. The fatty acid composition of breast milk is markedly altered based on maternal diet, which has important consequences for offspring as lipids are the principle energy source (up to 90%) for infants in early life (Delplanque *et al*., 2015). Like those exposed to maternal obesity, offspring exposed to a perinatal HFD (pHFD) are predisposed to developing metabolic disorders (Shrestha *et al*., 2020), anxiety and mood disorders (Rivera *et al*., 2015; DeCapo *et al*., 2019), as well as gastrointestinal (GI) dysfunction (Mulligan & Friedman, 2017; McMenamin *et al*., 2018; Clyburn *et al*., 2019) underscoring the importance of understanding the impact of diet itself on neural development.

Extrinsic control of the GI tract is mediated primarily by the parasympathetic nervous system via the vagus nerve. The cell bodies of vagal efferent motoneurons are contained within the dorsal motor nucleus of the vagus (DMV) which, together with the area postrema and the nucleus of the tractus solitarius (NTS), form the dorsal vagal complex (DVC) within the caudal hindbrain (Travagli & Anselmi, 2016). Vagal neurocircuits, in particular the NTS-DMV synapse, display a remarkable degree of neuroplasticity, with responsiveness varying depending on ongoing physiological conditions as well as environmental insults such as HFD or acute stress exposure (Browning & Travagli, 2014). Vagal neurocircuits develop relatively “late” with synaptic pruning and refinement occurring until at least postnatal day 28 (Rinaman & Levitt, 1993; Rao *et al*., 1997; Vincent & Tell, 1997). Exposure to a pHFD is known to decrease DMV neuronal excitability (decreased membrane resistance accompanied by a decreased ability to fire action potentials), alter DMV morphology (increased soma size and dendritic arborization) and prevent maturation of GABA_A_ receptor subunit structure and channel kinetics (retention of slower α2/3 GABA_A_ receptor subunits), all of which contribute to a decrease in basal gastric motility (Bhagat *et al*., 2015; Clyburn *et al*., 2019).

The perinatal period is also a time of increased vulnerability to stressors; early life stress (via maternal separation), for example, is associated delays in the development and maturation of descending hypothalamic-brainstem neurocircuits (Rinaman *et al*., 2011), while pHFD enables earlier activation of the hypothalamic-pituitary-adrenal (HPA) axis during the normally hyporesponsive early neonatal period (Abuaish *et al*., 2018), which may be contribute to the increased fear and anxiety responses seen in both non-human primates and rodents (Sasaki *et al*., 2013). How pHFD exposure affects the GI response to stress has not, however, been investigated.

As a response to an acute stress exposure, the body engages both the nervous, via the autonomic branches, and endocrine, via circulating hormones, systems to respond quickly to stress, leading to acute physiological changes at effector organs (Miller & O’Callaghan, 2002). Acute stress has two major effects on the motility of the GI tract; an increase in colonic transit (mediated by the sympathetic nervous system) (Stengel & Taché, 2009) and a decrease in gastric motility/tone with associated delays in gastric emptying rates (mediated by the vagus nerve) (Fone *et al*., 1990; Lewis *et al*., 2002; Nakade *et al*., 2005). In the CNS, both corticotropin releasing factor (CRF) and oxytocin (OXT) inputs from the paraventricular nucleus of the hypothalamus (PVN) project to the DVC and are critical to the ability of vagal neurocircuits to mount an appropriate gastric stress response (Mönnikes *et al*., 1992). CRF and OXT modulate DMV neuronal activity, both directly and indirectly, via modulation of NTS-DMV synaptic inputs (Dreifuss *et al*., 1992; Lewis *et al*., 2002; Holmes *et al*., 2013; Browning *et al*., 2014; Jiang *et al*., 2018; Jiang & Travagli, 2020).

Changes to hypothalamic innervation to vagal neurocircuits following pHFD exposure, and the effect this may have on their gastric physiology and stress response, has not been investigated.

The aim of the current study is to test the hypothesis that exposure to pHFD alters CRF and OXT hypothalamic inputs to the DMV, leading to dysregulated bagal brain-gut responses to stress

## Materials and Methods

### Ethical Approval

All experiments were performed with the approval of the Pennsylvania State University College of Medicine Institution Animal Care and Use Committee, and in accordance with the National Institute of Health regulations. Reporting of animal experiments conforms to the principles and regulations for animal experiment reporting and ethics and conforms with the ARRIVE guidelines.

### Animals

To generate the pHFD model, pregnant Sprague Dawley rats (Charles River, Kingston NY) were placed on either a control (fat, protein, and carbohydrate content 14:27:59% kcal; Purina Mills, Gray Summit, MO) or high fat (60:20:20% kcal; Envigo TD.06414) diet on embryonic day 13 (E13) to coincide with the *in utero* period when vagal neurocircuits begin to develop (Rinaman & Levitt, 1993; Rinaman *et al*., 2000) as well as to avoid induction of obesity in the dams. Dams remained on their respective diets through gestation and lactation. Litters were restricted to 12 pups after birth, and rat pups were weaned on postnatal day 21 (P21) onto their corresponding diets. Because of estrus-cycle dependent alterations in brainstem control of gastric functions and stress responses (Jiang *et al*., 2019), only male offspring (control N = 74; pHFD N = 72) were used in the current study. Electrophysiological studies were carried out on male rats that were at least 28 days of age, while in vivo studies (gastric motility recordings, gastric emptying assays) and neuronal tracing experiments were conducted on male rats that were at least 42 days of age. Rats were given access to food and water *ad libitum*.

In addition to the two diet groups, subsets of rats were also exposed to an acute stressor prior to gastric motility, gastric emptying, and electrophysiology experiments. Rats underwent a restraint stress (RS) which consisted of placing the rat in a cylinder (Plas-Labs; 544-RR) for 2hr ensuring their movement was restricted but caused no other physical duress. Fecal pellet output was then counted, and rats were used immediately in subsequent experiments.

### Neuronal Tracing experiments

At P46, 8 male rats (N=4/diet from 2 or more age-matched litters) were used for neuronal tracing experiments as described previously (Jiang *et al*., 2018). Briefly, rats received an injection of dexamethasone (Sigma D-6645; 1 mg/kg i.p.) to reduce brain swelling, before being anesthetized using a rodent cocktail (ketamine, xylazine, and acepromazine (80 mg/kg, 1.6 mg/kg, and 5 mg/kg, respectively, i.p.), Once a deep plane of anesthesia was reached, indicated by loss of the foot-pinch reflex, rats were placed on a stereotaxic frame (Kopf Instruments, Tujunga CA) on a homeothermic heating blanket to maintain body temperature of 37^0^C. The dorsal brainstem was exposed via blunt dissection and, following removal of the meningeal membrane, the vagal trigone was exposed. Then, five microinjections of Cholera Toxin B (CTB; List Labs, Campbell, CA, 0.5% in distilled water; 60nl per injection) were made using a glass pipette attached to a Picospritzer (Toohey Co., Fairfield NJ) into the rostro-caudal extent of the DMV at co-ordinates (in mm) rostro-caudal 0.0-0.6mm from calamus scriptorius, medio-lateral +0.3-0.4 from the midline; dorso-ventral −0.6-0.65 from the surface of the brainstem Injections were made over 2-3min and the pipette left in place for 5min between injections. The overlying musculature and skin were sutured (5/0 Vicryl suture) and rats returned to their home cages. Rats were given carprofen (Zoetis; 1 mg/kg, s.c.) for pain management for 3-5 days after surgery. After 12 days, the rats were anesthetized (Inactin, Sigma T133, 150mg/kg i.p.) and, once a deep plane of anesthesia was reached, perfused transcardially using 0.9% saline solution followed by 4% paraformaldehyde in PBS. Fixed brains were dissected, then postfixed and cryoprotected for 3 days in a 4% paraformaldehyde + 20% sucrose solution. Sections (50 micron-thick) throughout the entire extent of the hypothalamus and brainstem were collected using a freezing microtome, capturing the PVN and DVC, respectively. Tissue sections were kept at −20^0^C in a long-term storage buffer (50% 0.1M phosphate buffer (PBS; in mM: 115 NaCl, 75 Na_2_HPO_4_, and 7.5 KH_2_PO_4_):25% sucrose:25%ethylene glycol) until used for fluorescent immunohistochemistry.

Brainstem slices were used to visualize OXT and CRF fibers in the DVC as described previously (Jiang *et al*., 2018). Briefly, free floating slices were washed 3 times in PBS (0.1M), followed by incubation in Immunobuffer (0.3% Triton-X in TRIS-PBS) for 1 hour. Then, slices were incubated in donkey serum (1% in Immunobuffer; Abcam ab7475) for 1 hour to block nonselective binding. Slices were then incubated in goat anti-choline acetyl transferase (Millipore ab144p (ChAT); 1:1000) and rabbit anti-OXT (Sigma-Millipore ab911; 1:500) or rabbit anti-CRF (Abcam ab272391; 1:250) primary antibodies for 3 days at room temperature. Following repeated washing in PBS slices were incubated in donkey anti-goat IgG H&L (Invitrogen Alexa Fluor 488; 1:500) and donkey anti-rabbit IgG H&L (Alexa Fluor 568; 1:500) secondary antibodies overnight at room temperature. Hypothalamic slices were visualized using the same protocol staining with goat anti-cholera toxin B (List Labs 703 (CTB): 1:1000) and either rabbit anti-OXT (1:500) or rabbit anti-CRF (1:250) primary antibodies. After washing in PBS, brain slices were mounted on subbed slides (Superfrost Microscope Slides; Fisher Scientific) and cover-slipped using Fluoromount-G (Invitrogen).

### Immunohistochemical measurements

Immunohistochemical tissues were examined using a Zeiss LSM 900 compact microscope. Confocal images throughout the depth of the 50 micron-thick slices were taken at 20X magnification and compiled to a z-stack. An image was taken of each side (left and right) of the DVC and the PVN.

Sequential scanning and adjustment of the appropriate High-Low settings allowed for optimal intensities. All images were taken at the same settings to allow for direct comparison between samples. The intensity of OXT and CRF fibers within the DMV or NTS was analyzed using ImageJ software (https://imagej.nih.gov/ij/). To determine the intensity of fibers, the DMV or NTS were outlined and background staining was normalized, then mean pixel intensity was calculated. Mean pixel density was calculated in both the left and right side. For each rat, an image was taken at the intermediate (Bregma −13.9mm) level of the brainstem. The number of CRF+, OXT+, and CTB+ neurons in the PVN was counted using ImageJ software. Cell counts were made in both the left and right side of and throughout the length of the PVN, noted as anterior, intermediate, or posterior based on anatomical landmarks (Bregma levels −0.92, −1.80, and −2.30mm, respectively). Co-localization of signal was identified by cells with white fluorescence and counted as such.

### Protein measurement

Male rats from both diet groups and stress conditions (N=10/group from 2 or more age-matched litters) were randomly selected for serum extraction. Rats were anesthetized with isoflurane (5% in air) and, once a deep plane of anesthesia was induced, a cardiac puncture was performed to obtain truncal blood samples. A small amount of blood was used to measure blood glucose levels using a glucometer (One Touch Ultra meter) and reported as mg/dL. The remaining blood was left to clot for approximately 60min at room temperature before serum was extracted after centrifugation (Eppendorf Centrifuge 5417R; 1500 rpm for 4 minutes). Samples were stored at −20°C until use. Serum samples were run in duplicate using the Enzo Corticosterone ELISA kit (Farmingdale, NY, USA) at a 1:40 dilution (26.99 pg ml^−1^ sensitivity). The optical density of the samples was measured using a SpectraMax 340PC 384 plate reader interfaced with SoftMaxPRO software at 405nm. The calibration concentration curve was calculated using an online freely-available immunoassay software package utilizing a 4-parameter logistic curve fitting program (https://www.myassays.com), and unknown concentrations were determined via interpolation.

Adult male rats from both diet groups (N=17-23 rats/diet from 2 or more age-matched litters) were randomly selected for cerebrospinal fluid (CSF) extraction. Rats were anesthetized with Inactin (Sigma T133; 150mg/kg i.p.). Once anesthetized deeply (loss of foot-pinch withdrawal reflex), rats were placed on a stereotaxic frame and the dorsal brainstem was exposed via blunt dissection. At this time, 100-200µL of CSF were extracted using a pipette over a period of approximately 5 minutes. Directly after, the sample was centrifuged (Eppendorf Centrifuge 5417R; 1500 rpm for 4 minutes) and the freshly separated CSF was immediately stored at −80°C until use. CSF samples from control (N=10) and pHFD (N=8) rats were run using the Enzo Oxytocin ELISA kit (Farmingdale, NY, USA) according to manufacture instructions (15.0 pg ml^−1^ sensitivity). The optical density of the samples was measured using a SpectraMax 340PC 384 plate reader interfaced with SoftMaxPRO software at 405nm. The calibration concentration curve was calculated using an online freely-available immunoassay software package utilizing a 4-parameter logistic curve fitting program (https://www.myassays.com), and unknown concentrations were determined via interpolation. CSF samples from control (N=13) and pHFD (N=9) rats were also run using the Biomatik CRF ELISA Kit (Kitchener, Ontario, Canada) and according to manufacture instructions were run neat (5.26 pg ml^−1^ sensitivity). The optical density of the samples was measured using a SpectraMax 340PC 384 plate reader interfaced with SoftMaxPRO software at 450nm. The calibrator concentration curve and unknown concentrations were calculated as the other ELISA experiments.

### Electrophysiology

Electrophysiological recordings were made from unstressed and stressed (2hr RS) control and pHFD rats that were at least 28 days old. Rats were anesthetized with isoflurane (5% in air) before euthanized via administration of a bilateral pneumothorax. The brainstem was removed and sliced as previously described (Clyburn *et al*., 2021). Briefly, the brainstems were excised quickly and submerged rapidly in cold (4°C) oxygenated Krebs’ solution before being mounted on a vibratome and 300µm sections throughout the rostro-caudal extent of the brainstem were cut. Slices were placed in warm (30°C) oxygenated Krebs solution for 90min before recording. For each recording, the brainstem slice was placed in the perfusion chamber on the stage of a Nikon E600FN microscope. Slices were perfused with warmed Krebs solution (in mM: 126 NaCl, 25 NaHCO_3_, 2.5 KCl, 1.2 MgCl_2_, 2.4 CaCl_2_, 1.2 NaH_2_PO_4_, and 10 D-glucose kept at pH 7.4 by bubbling with 95% O_2_-5% CO_2_) maintained at 32°C at a rate of 2.0 –2.5 ml/min. DMV neurons were identified by their size and relative location to the dorsal NTS and ventral hypoglossal nucleus. Electrophysiological recordings of 79 DMV neurons (N=16-26/group with recordings from a minimum of 3 rats from at least 2 litters/experiment) were made under bright-field illumination using 2- to 4-MΩ patch pipettes filled with a potassium chloride intracellular solution (in mM: 140 KCl, 1 CaCl_2_, 1 MgCl_2_, 10 HEPES, 10 EGTA, 2 NaATP, and 0.25 NaGTP adjusted to pH 7.4) and a single electrode voltage-clamp amplifier (Axopatch 200B; Molecular Devices). Data were filtered at 2 kHz, digitized via a Digidata 1440 Interface, and stored an analyzed on a PC with pClamp 10 software (Molecular Devices). Recordings with a series resistance larger than 20MΩ were eliminated from the study.

Miniature inhibitory postsynaptic currents (mIPSCs) were measured in the voltage-clamp configuration, at a holding potential of −50 mV. mIPSCs were isolated by perfusing the cell with tetrodotoxin (TTX; 1µM) and kynurenic acid (KynA; 1mM), a glutamatergic ionic receptor antagonist, to isolate inhibitory action potential-independent neurotransmitter release. All drugs were made up in fresh Krebs’ solution on the day of the experiment. Baseline recordings were taken after at least 5min of TTX + KynA application, followed by application of either OXT (Bachem 4016373; 100nM) or the CRF1 receptor antagonist, astressin (Bachem 4026664; 1µM). Miniature current characteristics, including changes in amplitude (pA), frequency (Hz), and charge transfer (frequency (Hz) * area (pAms)), were assessed using the Mini Analysis Program (Synaptosoft).

### Gastric motility recordings

A subset of 18 rats (N=9/diet from 2 or more age-matched litters) were randomly chosen to undergo gastric motility recordings as described previously (Clyburn *et al*., 2021). Briefly, rats were fasted for 18 hours before experimentation and anesthetized with Inactin (Sigma T133; 150mg/kg i.p.). A tracheotomy was performed prior to making a midline abdominal incision to expose the ventral stomach. Then, a strain gauge (MSR Neurobiology, Modern Scientific Research LLC, Roaring Spring, PA) was fixed to the gastric corpus, running parallel to the border between corpus and fundus. Rats were placed on a stereotaxic frame and the dorsal brainstem was exposed via blunt dissection. Then, the meningeal membrane was removed and the vagal trigone was exposed. Rats were left to rest for at least 1 hour, allowing for recovery from surgical manipulation. Recovery was noted as consistent gastric contractile activity with a stable baseline. Recordings of the corpus contractile activity was filtered (MSR Neurobiology), amplified (QuantaMetrics EXP CLSG-2), then visualized using Axoscope software (Molecular Devices). To ensure proper placement of the microinjection pipette, bicuculline (BIC; 25pmols/60nl) was first injected into the left DMV at coordinates (in mm) +0.2 to 0.3 rostro-caudal from calamus scriptorus, −0.2 to 0.3 medio-lateral from the midline, and −0.6 to 0.65 dorsal-ventral from the surface of the brainstem (60nl over 1min). Once an increase in gastric tone and motility was observed following BIC injection, these coordinates were used for subsequent injections.

Once the stereotaxic coordinates of the DMV were confirmed, baseline corpus motility was recorded for at least 5 min before oxytocin (OXT; 150 pmols/60nl) was microinjected with recording continuing until baseline was restored with at least 1 hour being allowed between microinjections. After recording another baseline period of activity, astressin (10µg/2µl) was applied to the 4^th^ ventricle using a pipette, with two doses given at 5min intervals, prior to repeated OXT microinjection. Changes to corpus tone and motility were measured as described previously (Clyburn *et al*., 2021). Briefly, strain gauges were calibrated prior to use and drug-induced effects on gastric motility and tone were extrapolated from the average calibration value. While basal gastric tone was not adjusted to a fixed value, the gastric circular muscle provided a basal tension of approximately 500mg. Due to individual variation in animal size and surgical placement of the strain gauges which may lead to minor variations in absolute values, corpus tone is reported as absolute values relative to baseline. Each rat served as its own control and corpus motility was calculated using the following formula:

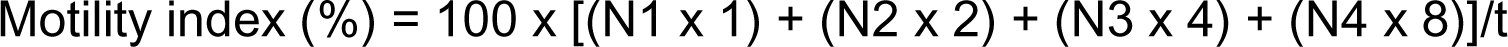

Where Nx = the number of motility peaks in each force range and t = time over which motility was measured (seconds). With the assumption that the absence of motility produced a 0mV signal, the peak-to-peak motility waves reflected N1 = 25-50mg, N2 = 51-100mg, N3 = 101-200mg, and N4 = >201mg. At the completion of the experiment, the rat was perfused with 0.9% saline followed by 4% PFA. The brainstem was dissected, and 50μm thick sections made as described previously to confirm the site of injection.

### Gastric Emptying

Male rats (N=8/diet from 2 or more age-matched litters) were randomly selected to undergo the ^13^C gastric emptying assay, at 6 weeks of age, to determine the emptying rate of solid food as described previously (Clyburn *et al*., 2021). Briefly, rats were first acclimated to the testing procedure by being placed in their respective holding chamber (VetEquip, Livermore CA) for approximately 3 hours/d for 3 consecutive days. While in the chamber, they were fed 1g of pancake (De Waffelbakkers Buttermilk Pancakes). After the 3rd acclimation day, rats were fasted overnight, with *ad libitum* access to water, prior to testing. On test day, rats were placed in their holding chamber for 1hr to allow the carbon dioxide isotope analyzer (Los Gatos Research, San Jose CA) to equilibrate. Then, each rat was fed 1g of a pancake that contained 4 µl of [^13^C]-octanoic acid (Cambridge Isotope Laboratories, Inc, Tewksbury MA). After visually confirming the rats had eaten the entire pancake (in less than 5min or they were omitted from the study), the rats were left to remain in their chamber for at least 4hr. During this time, air from each of the testing chambers was automatically sampled, one at a time, at 30s intervals. At the completion of the gastric emptying study, rats were returned to their cages with full access to food and water. Rats were re-acclimated to their testing chambers 3-4d after testing, and gastric emptying was repeated weekly for 3 baseline measurements. The following week, rats underwent the 2hr restraint stress protocol before being placed in the chambers for data collection.

To analyze these data, the first 10s of each 30s period of data were removed to ensure a complete flush of the previous air sample from the tubing. The remaining 20s of data were averaged for a single data point at a given time. The concentration of ^13^CO_2_ was then calculated and expressed as a change over the baseline. The change of concentration of ^13^CO_2_ vs time (t) was fitted by a non-linear regression curve using SigmaPlot with the following equation (y = at^b^e^-ct^) where y is the percentage of the ^13^C excretion in the breath per hour (t) and a, b, and c are regression constants estimated for each breath vs. time curve. The gastric half emptying time (T_1/2_) was calculated from a numerical integration procedure using an inverse gamma function. The three baseline T_1/2_ values for each animal were averaged.

### Statistical analyses

Statistical analysis was conducted using Prism (GraphPad, Boston MA) software. Results are expressed as mean ± SD of the indicated sample size (N; cells or rats) with statistical significance defined as P<0.05.

For neuronal tracing experiments, PVN neurons were identified by immunofluorescence with a recognizable nucleus. Co-staining was counted with the appearance of white fluorescence signal. Neuronal cell counts were compared between diet groups (control and pHFD) using a two-way unpaired t-test. Immunofluorescent fibers within the brainstem were quantified as a fluorescent area (thresholded using Image J software) expressed as a percentage of the total brain region identified and compared between diet groups (control and pHFD) using a two-tailed unpaired t-test.

For experiments measuring corticosterone serum protein levels, protein concentration is reported as pg/mL based on optical density, and calculated via a concentration curve generated using MyAssay software. Protein concentrations were grouped by diet (control or pHFD) and stress (unstressed or stressed), and compared using a two-way ANOVA with post-hoc Bonferroni test to identify specific group differences.

For experiments measuring CSF OXT and CRF, protein concentration is reported as pg/mL based on optical density, and calculated via a concentration curve generated using MyAssay software. Concentration values of the specified protein were grouped by diet (control or pHFD). Groups were compared using a two-tailed unpaired t-test.

When analyzing gastric motility and tone, each rat served as its own control with corpus responses (either changes in motility or tone) being assessed and compared before and after drug application (either direct microinjection into the DMV or 4^th^ ventricular application) using a two-tailed paired t-test. Inter-group comparisons (control vs. pHFD exposure) were made using a two-tailed unpaired t-test. Only rats that responded to bicuculline with an increase in tone and motility, to ensure proper placement of the microinjection pipette, were included in the statistical analysis.

Gastric emptying is expressed as time (min) or as a change from baseline (min), with baseline defined as the average half-emptying (T_1/2_) time of 3 baseline measurements (taken over the course of 3 weeks). Baseline gastric emptying rates were grouped based on diet (control or pHFD) and compared using a two-tailed unpaired t-test. Gastric emptying rates after an acute stressor were compared within each rat (average baseline rate vs. rate after stressor) using a two-tailed paired t-test. The magnitude of change after a stressor between diet groups (represented as a change in time (min); control vs. pHFD) was compared using a two-tailed unpaired t-test.

For electrophysiological results, the magnitude of the response of any neuron to drug application was assessed using one sample t-test against a theoretical mean of 100% (indicating no change from baseline). The magnitude of the drug response was determined by calculating the difference between baseline and drug conditions and transformed into a percentage of baseline. A change from baseline greater than 25% indicates a physiologically significant change. Any neuron that had a baseline frequency of less than 0.5 Hz was not included in the analysis. The magnitude of response to an individual drug was compared within control diet groups (stressed and unstressed), pHFD groups (stressed and unstressed), and between unstressed groups (control vs pHFD) using a two-tailed unpaired t-test.

### Chemicals and Drugs

Oxytocin and astressin were purchased from Bachem (Torrance, CA), TTX was purchased from Cayman Chemical Company (Ann Arbor, MI); all other chemicals were purchased from Millipore Sigma (St. Louis, MO).

## Results

### Acute stress exposure increases fecal pellet output, blood glucose, and serum corticosterone levels in control and pHFD rats

Age-matched rats in both control (N=30) and pHFD (N=31) diet groups gained weight in a similar manner (P_Time_ <0.0001), however the weights in both groups did not differ at each time point studied (P_Diet_ =0.2342, using a mixed-effects model; Figure 1A). These results indicate that the pHFD rats were not yet obese, validating the postulate that our experimental model analyses the effect of diet specifically.

**FIGURE 1:**
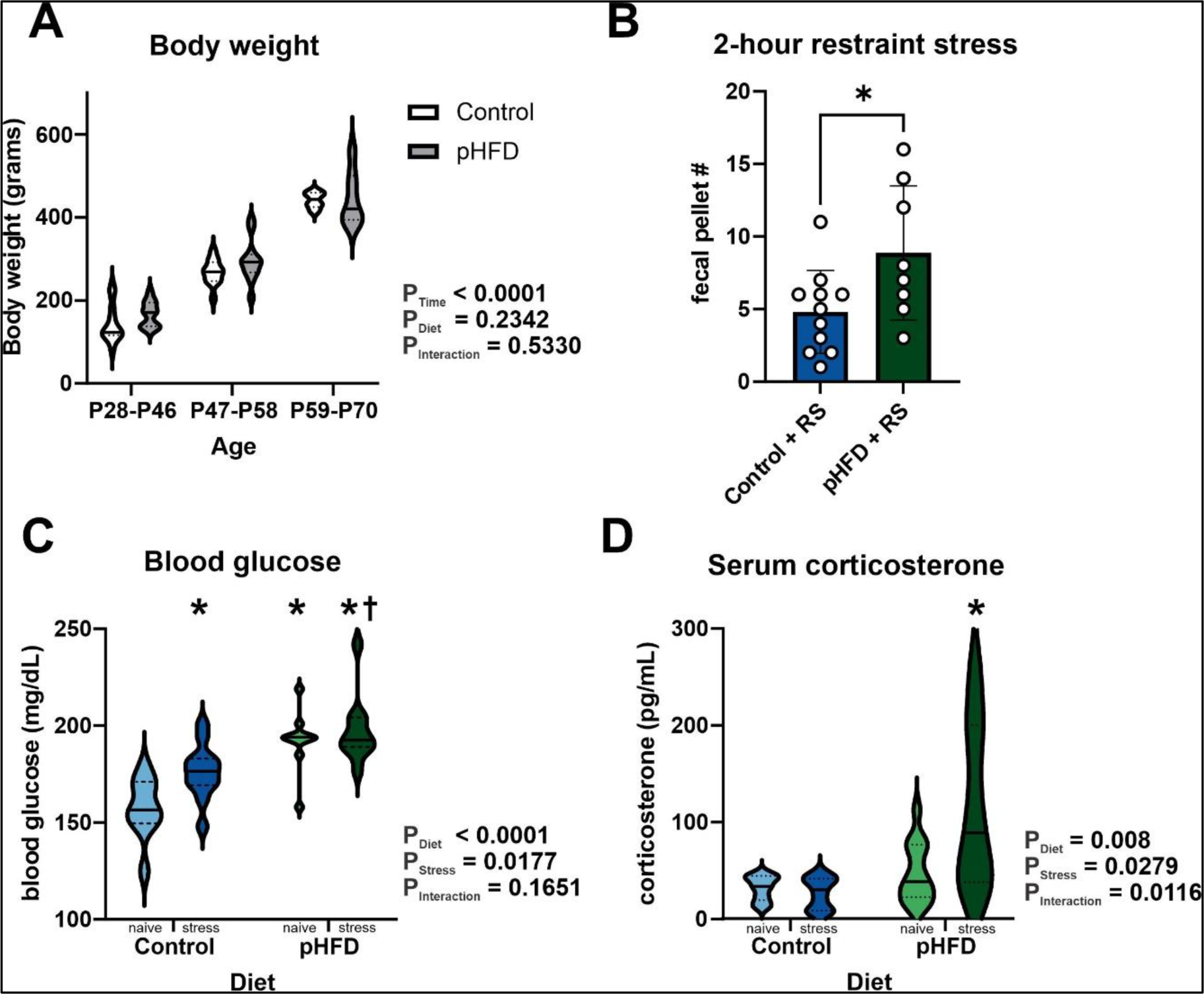
pHFD exposure increases fecal pellet output and elevates blood glucose levels in response to acute stress. **A**: Graphical summary of body weights (g) in control (N= 6-12 per group age) and pHFD (N= 7-12 per group age) rats across experimental timepoints; P28-P46, P47-P58, and P56-P70. Data demonstrated time, but not diet, increased body weight and pHFD rats were neither overweight norobese, compared to age-matched control rats (Mixed-effects model with REML estimation; p<0.05 for significant effect). **B:** Graphical summary of fecal pellet output during 2-hour RS showing a significant increase in fecal pellet output in control (N=11) and in pHFD (N=8) rats (* p<0.05 using two-way unpaired t-test). **C:** Graphical summary of blood glucose levels (mg/dL) of control (N=10, light blue), control + RS (N=12, dark blue), pHFD (N=10, light green), and pHFD + RS (N=10, dark green) at time of tissue collection. A two-way ANOVA analysis, with diet and stress as variables, revealed that both affected blood glucose levels but that they did not interact (p<0.05 denotes a significant effect). *Post-hoc* Bonferroni multiple comparisons test indicated that blood glucose levels were significantly elevated in control + RS and both pHFD groups compared to unstressed controls (* p<0.05), demonstrating that stress and exposure to a pHFD elevates blood glucose levels. Additionally, blood glucose levels were elevated in pHFD + RS rats compared to control + RS rats († p<0.05) indicating that pHFD exposure exacerbated the stress-induced response. **D:** Graphical summary of serum corticosterone levels (pg/mL) of control (N=10, light blue), control + RS (N=9, dark blue), pHFD (N=13, light green), and pHFD + RS (N=14, dark green) rats at time of tissue collection showing significant effects of diet, stress, and their interaction (p<0.05 denotes significance; two-way ANOVA). While corticosterone levels returned to unstressed levels in control animals after 2-hour RS stress, they remained significantly elevated in pHFD rats (* p<0.05 compared to all groups; *post-hoc* Bonferroni multiple comparisons test).

A two-hour restraint stress (RS) model was chosen as it is a moderate psychological stressor (Buynitsky & Mostofsky, 2009) shown previously to induce reliable changes to gastric physiology (Nakade *et al*., 2005; Zheng *et al*., 2009; Jiang *et al*., 2018; Jiang *et al*., 2019). pHFD rats (N=8) had a significantly larger fecal pellet output, a validated indicator of stress, in response to the restraint stress than control (N=11) rats (8.8±4.61 fecal pellets vs 4.8±2.86 fecal pellets in stressed pHFD vs control rats, respectively; p=0.0298, t(17)=2.371, 95% C.I.=0.4469 to 7.667; two-tailed unpaired t-test; Figure 1B) suggesting that the colonic response of pHFD rats is more sensitive to RS.

Elevations in blood glucose levels is a peripheral marker of stress (Christiansen *et al*., 2007). Blood collection was made at the time of sacrifice in control (N=10), control + RS (N=12), pHFD (N=10), and pHFD + RS (N=10) rats, with results indicating blood glucose levels were significantly increased by both stress (F(1, 38)=6.154, P_Stress_ =0.0177) and diet (F(1, 38)=34.52, P_Diet_ <0.0001; two-way ANOVA; Figure 1C). In control rats, stress elevated blood glucose levels significantly (from 157.6 to 176.3 mg/dL vs; p=0.0459, t(38)=2.817, 95% C.I.=-37.08 to −0.2214; *post-hoc* Bonferroni multiple comparison test; Figure 1C). pHFD rats had significantly elevated blood glucose levels compared to controls (192.5 vs 157.6 mg/dL in pHFD vs control rats, respectively; p<0.0001, t(38)=5.047, 95% C.I.=-54.15 to −15.65; Figure 1C) and pHFD + RS rats had significantly elevated blood glucose levels compared to control + RS rats (197.6 vs 176.3 mg/dL in stressed pHFD vs stressed control rats, respectively; p=0.0156, t(38)=3.225, 95% C.I.=-39.78 to −2.921; Figure 1C). RS itself did not, however, increase blood glucose levels significantly in pHFD rats (197.6 vs 192.5 mg/dL in stressed pHFD vs pHFD rats, respectively; p=0.9766, t(38)=0.7375, 95% C.I.=-24.29 to 14.09; Figure 1C).

These results indicate that pHFD rats have elevated basal levels of blood glucose, but they do not mount a hyperglycemic response to stress like control animals.

An increase in plasma corticosterone (CORT) levels after stress exposure is a suitable marker of HPA activation and general stress reactivity (Leistner & Menke, 2020). Serum CORT levels were collected at the time of sacrifice in control (N=10), control + RS (N=9), pHFD (N=13), and pHFD + RS (N=14) rats. The data indicate that stress, diet, and their interaction increased CORT levels significantly (F(1, 44)=4.726, P_Stress_ =0.0351; F(1, 44)=12.93, P_Diet_ =0.0008; F(1, 44)=6.384, P_Interaction_ =0.0152; two-way ANOVA; Figure 1D).

CORT levels were not found to be different between the control and pHFD unstressed groups (32.26 vs 47.67 pg/mL; p>0.9999, t(44)=0.7789, 95% C.I.=-70.11 to 39.27; post-hoc Bonferroni multiple comparison test) but were significantly more elevated in stressed pHFD rats compared to stressed control rats (115.5 vs 27.16 pg/mL; p=0.0008, t(44)=4.208, 95% C.I.=-146.3 to −30.33; Figure 1D). Furthermore, stresssignificantly elevated CORT levels in pHFD rats (115.5 pg/mL vs 47.67 pg/mL; p=0.0041, t(44)=3.652, 95% C.I.=-119.1 to −16.51Figure 1D), but not in control rats (27.16 pg/mL vs 32.26 pg/mL; p>0.9999, t(44)=0.2305, 95% C.I.=-55.92 to 66.10).

Taken together, they data indicate that while basal CORT levels did not differ between diet groups, stress causes a significant and prolonged elevation in pHFD rats.

### pHFD slows basal gastric emptying

Previous studies have shown that pHFD decreases basal gastric tone and motility in a vagally-dependent manner (Clyburn *et al*., 2019). Gastric emptying rates were assessed in control (N=8) and pHFD (N=8) male rats using the ^13^C octanoic acid breath test. The rate of gastric emptying in pHFD rats was significantly delayed compared to control rats (76.9±22.07 minutes vs. 57.6±6.94 minutes, p=0.0338, t(14)=2.352, 95% C.I.=-36.79 to −1.699; two-tailed unpaired t-test; Figure 2B).

**FIGURE 2:**
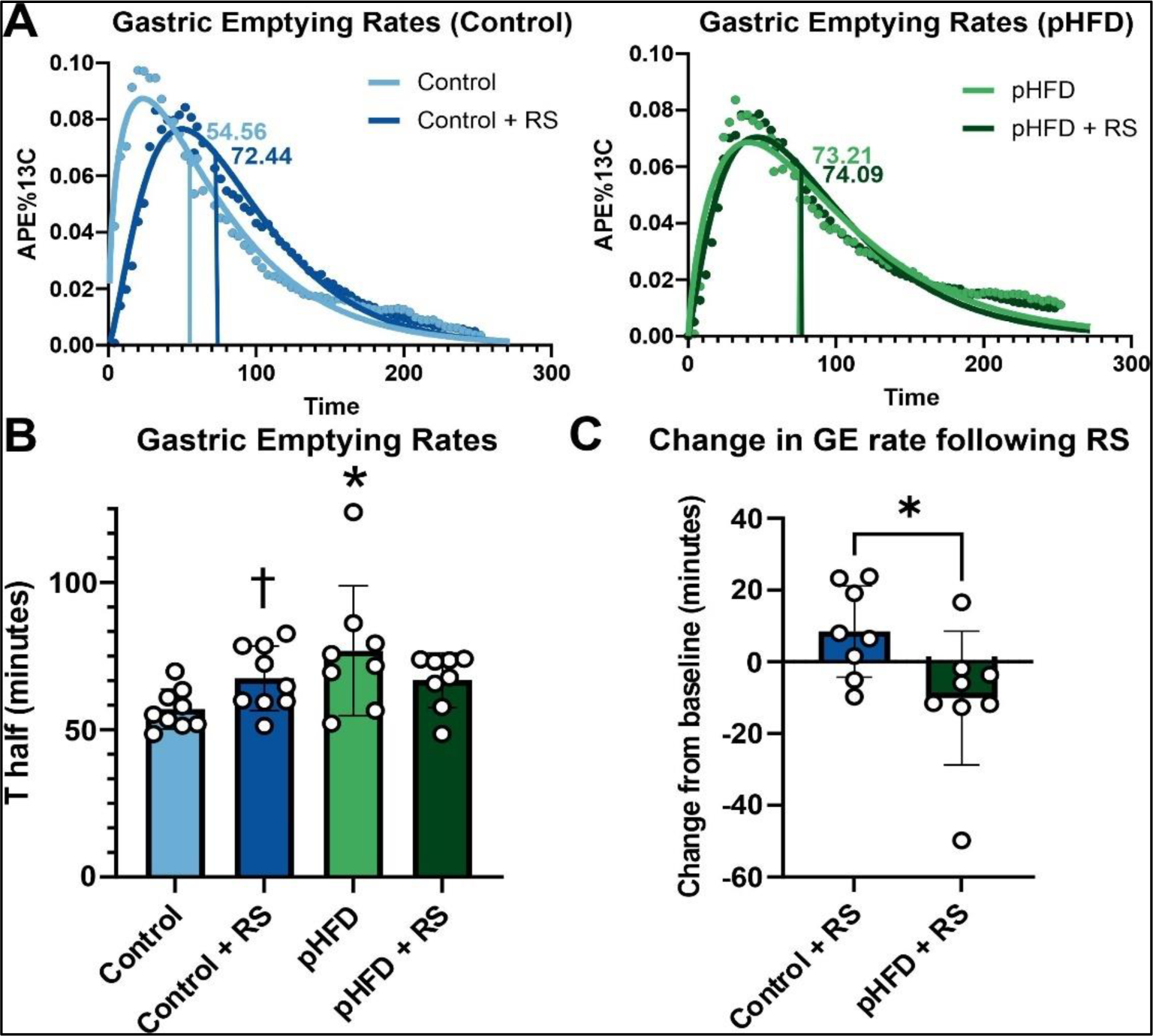
pHFD rats have slower basal gastric emptying rates and do respond to stress with a further delay in gastric emptying. **A:** Representative gastric emptying curves of control (left; light blue), control + RS (dark blue), pHFD (right; light green), and pHFD + RS (dark green) rats. Vertical lines indicate the T_1/2_ values. RS delayed gastric emptying rates (seen by an increase in T_1/2_ value) in control, but not pHFD, rats. **B:** Graphical summary of gastric emptying rates in control (N=9) and pHFD rats (N=8) before and after stress exposure. Data indicate that pHFD rats had a significantly slower basal gastric emptying rates vs control rats (* p<0.05, two-tailed unpaired t-test comparing unstressed control to unstressed pHFD T_1/2_ values) and that 2-hour RS significantly delayed gastric emptying in control, but not pHFD, rats († p<0.05, two-tailed paired t-test comparing unstressed and stressed T_1/2_ values among diet groups). **C:** Graphical summary of the change in gastric emptying rates (minutes) after stress (stressed-unstressed) demonstrating the significant difference in response between both diet groups (* p<0.05, two-tailed unpaired t-test comparing the change in T_1/2_ values).

Acute stress is known to decrease gastric motility and tone as well as delay gastric emptying (Taché & Perdue, 2004; Bhatia & Tandon, 2005; Jiang & Travagli, 2020). To assess whether pHFD dysregulates the response to acute stress, gastric emptying was assessed in rats before and after 2hr RS. As shown previously (Babygirija *et al*., 2010; Jiang & Travagli, 2020), control rats responded to acute stress with a delay in gastric emptying (from 57.0±6.765 to 67.5±10.92 minutes in naïve and stressed rats respectively, p=0.0466, t(8)=2.351, 95% C.I.=0.2013 to 20.73; two-tailed paired t-test). In contrast, pHFD rats showed no significant change in gastric emptying rates after stress (76.9±22.07 and 66.8±9.30 minute, in naïve and stressed rats respectively; p=0.1703, t(14)=2.321, 95% C.I.=-35.61 to −1.404; two-tailed paired t-test; Figure 2A,B). Comparing the change in gastric emptying rates following RS between diet groups revels a significant difference in the change of emptying rate from the unstressed condition (+8.443±12.72 minutes vs −10.06±18.62 minutes; p=0.0359, t(14)=2.321. 95% C.I.=-35.61 to −1.404; two-tailed unpaired t-test; Figure 2C).

Taken together, these data demonstrate that pHFD exposure results in a slower basal gastric emptying rate and prevents the appropriate delay in gastric emptying in response to an acute stress.

### pHFD decreases PVN^OXT^ DMV-projecting neurons and OXT fibers in the DMV

### pHFD increases PVN^CRF-^DMV neurons but decreases CRF fibers in the DMV

The PVN provides the only source of OXT innervation to the DVC, and OXT modulates gastric functions, particularly in response to stress (Jiang *et al*., 2018; Jiang & Travagli, 2020). The number of PVN^OXT^ neurons projecting to the DMV were assessed following DMV microinjection of the retrograde tracer, CTB, in control and pHFD rats (N=4/diet). Immunofluorescent images of anterior, intermediate, and posterior sections of the PVN (Bregma levels −0.92, −1.80, and −2.30mm, respectively) were visualized to allow assessment of DMV-projecting CTB+ neurons, OXT+ neurons, and CTB+/OXT+ on each side of each PVN section. Following pHFD, fewer PVN-DMV neurons were identified in the intermediate PVN (80.1±26.09 vs 149.8±61.4 cells/section in pHFD and control rats, respectively; p=0.0019, t(18)=3.640, 95% C.I.=-109.9 to −29.46; two-tailed unpaired t-test; Figure 3G), although no significant differences were found in the number of PVN^OXT^-DMV neurons in the anterior (32.1±36.65 vs. 60.2±32.36 cells/section in pHFD and control rats, respectively, p=0.1079. t(16)=1.703, 95% C.I.=-63.18 to 6.885; two-tailed unpaired t-test, Figure 3D) or posterior (120.4±55.97 vs. 97.2±72.95 cells/section in pHFD and control rats, respectively, p=0.5120, t(12)=0.6757, 95% C.I.=-51.63 to 98.05; two-tailed unpaired t-test, Figure 3J) PVN.

**FIGURE 3:**
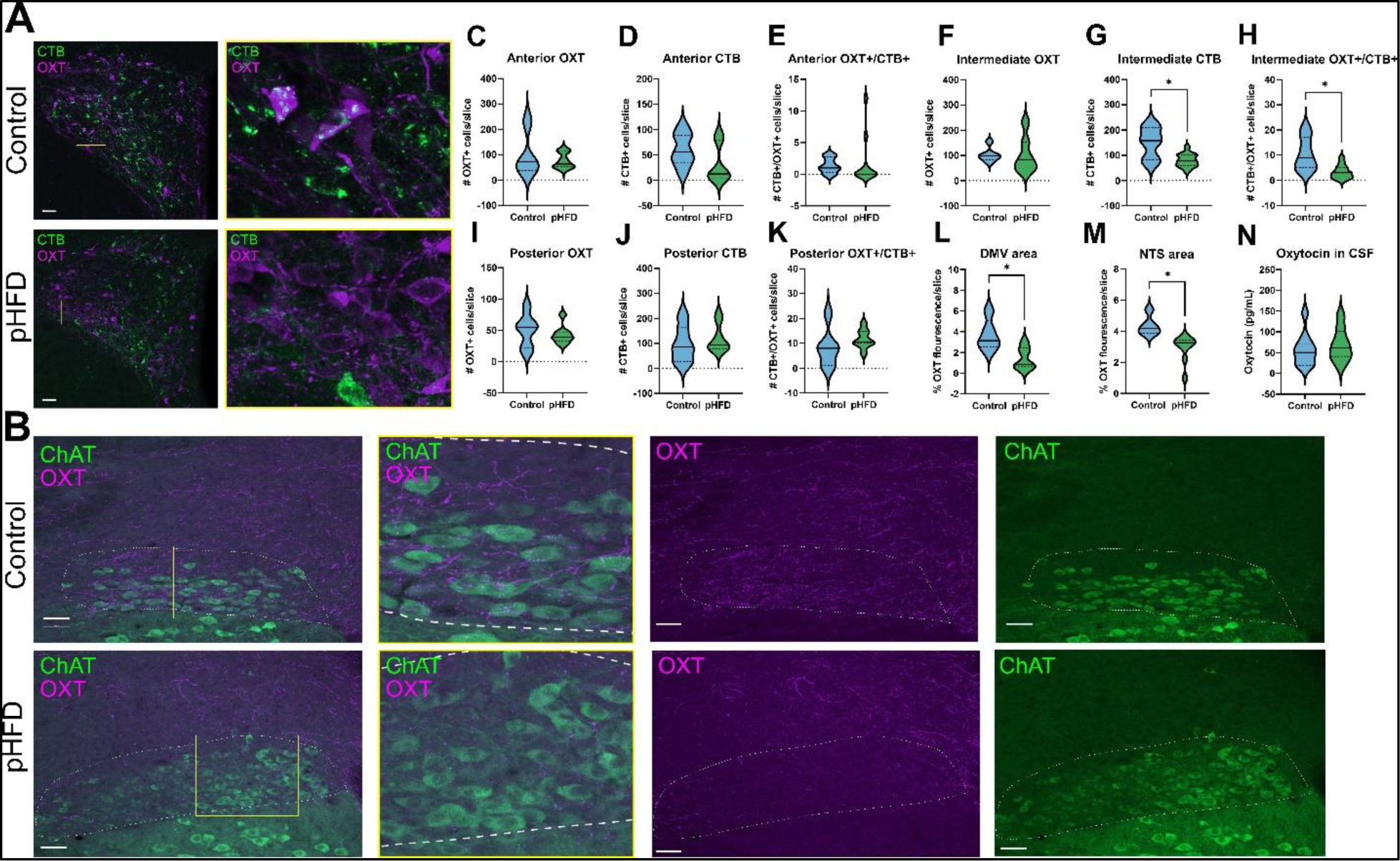
pHFD decreases DMV-projecting PVN^OXT^ neurons and reduces OXT fiber expression in the DMV and NTS. **A:** Representative images of the intermediate PVN with CTB+ neurons in green and OXT+ neurons in magenta from a control (top) and pHFD (bottom) rat, illustrating the loss of CTB+ and CTB/OXT co-localized neurons following pHFD exposure. Scale bar represents 50 µm. pHFD (N=4) and control (N=4) rats received a microinjection of the retrograde tracer CTB in the DMV. **B:** Representative images of the brainstem containing the DMV (outlined; visualized by ChAT-IR to identify cholinergic neurons) and NTS in control (top) and pHFD (bottom) rats, illustrating the loss of OXT fibers following pHFD exposure. Scale bar represents 50 µm. Quantification of neurons in the anterior PVN found no differences in the number of OXT+ (**C**), CTB + (**D**), and OXT+/CTB+ (**E**) cells between diet groups (N=4 for each). **F:** Graphical summary of quantification of OXT+ cells in the intermediate PVN, showing no difference between control and pHFD rats. **G:** Graphical summary of quantification of CTB+ neurons in the intermediate PVN, showing there was a significant reduction in DMV-projecting neurons in pHFD rats (* p<0.05 using a two-tailed unpaired t-test). **H:** Graphical summary of quantification of OXT and CTB co-localization showing a significant reduction of DMV-projecting PVN^OXT^ neurons in pHFD rats (* p <0.05 using two-tailed unpaired t-test). Quantification of neurons in the posterior PVN found no differences in the number of OXT+ (**I**), CTB+ (**J**), and OXT+/CTB+ (**K**) cells between diet groups. Graphical summary of % OXT fluorescence in the DMV (**L**) and NTS (**M**) illustrating the significant reduction in pHFD compared to control rats (* p<0.05 for each using a two-tailed unpaired t-test). **N:** Graphical summary of CSF OXT levels in pHFD (N=8) and control (N=10) rats demonstrating no significant difference between the diet groups.

The results also showed a reduction in specifically PVN^OXT-^DMV neurons (3.3±2.7 vs 10.5±6.6 cells/section; p=0.0032 in pHFD and control rats, respectively, t(18)=3.405, 95% C.I.=-11.59 to −2.745; two-tailed unpaired t-test; Figure 3H) in the intermediate PVN, with no significant difference in anterior PVN^OXT-^DMV neurons (1.900±4.012 vs. 1.375±1.188 cells/section in pHFD and control rats, respectively, p=0.7266, t(16)=0.3559, 95% C.I.=-2.603 to 3.653; two-tailed unpaired t-test, Figure 3E) or posterior PVN^OXT-^DMV neurons (11.75±3.732 vs. 8.167±7.679 cells/section in pHFD and control rats, respectively, p=0.2685, t(12)=1.160, 95% C.I.=-3.145 to 10.31; two-tailed unpaired t-test, Figure 3K).

In contrast, pHFD did not alter the overall number of PVN^OXT^ neurons at either the intermediate (102.9±68.7 vs. 102.3±28.37 cells/section in pHFD and control rats, respectively, p=0.9843, t(17)=0.02001, 95% C.I.=-61.59 to 62.77; two-tailed unpaired t-test; Figure 3F), anterior (75.60±30.03 vs. 98.50±84.61 cells/section in pHFD and control rats, respectively, p=0.4353, t(16)=0.8003, 95% C.I.=-83.56 to 37.76; two-tailed unpaired t-test, Figure 3C), or posterior (42.75±14.77 vs. 49.50±26.20 cells/section in pHFD and control rats, respectively, p=0.5501, t(12)=0.6148, 95% C.I.=-30.67 to 17.17; two-tailed unpaired t-test, Figure 3I) level of the PVN, suggesting that effects of pHFD were selective for those PVN^OXT^ neurons innervating the DMV.

Immunohistochemistry was used to quantify OXT fibers at the intermediate level of the DVC (Bregma level −7.34) in both the DMV and NTS. ChAT-immunoreactivity (-IR) was used to identify cholinergic vagal motoneurons and delineate the boundaries of the DMV and the fluorescence intensity of the OXT-IR was quantified in both sides of the DMV and NTS using Image J. pHFD significantly reduced OXT-IR in the NTS (2.937±0.849% vs 4.345±0.628% fluorescence intensity in pHFD and control rats, respectively, p=0.0073, t(12)=3.225, 95% C.I.=-2.359 to −0.4568; two-tailed unpaired t-test; Figure 3M), and DMV (1.431±1.061% vs 3.672±1.494% fluorescence intensity/area in pHFD and control rats, respectively, p=0.0121, t(10)=3.057, 95% C.I.=-3.875 to −0.6079; via two-tailed unpaired t-test; Figure 3L).

Lastly, since previous studies have demonstrated that OXT within the CSF may arise from the PVN (Martínez-Lorenzana *et al*., 2008), CSF samples from the fourth ventricle were taken from control and pHFD rats. The results showed no significant difference in basal OXT levels in the CSF between control and pHFD rats (56.7±38.81 vs 68.8±39.66 pg/ml in pHFD and control rats, respectively; p=0.5223, t(16)=0.6542, 95% C.I.=-27.24 to 51.56; two-tailed unpaired t-test, Figure 3N). These data support the overall lack of change in PVN OXT neuronal numbers following pHFD.

Taken together, these results indicate that pHFD exposure leads to a selective reduction in OXT neurons that project to the DMV where there are reduced OXT fibers.

The PVN also sends CRF projections to the DVC, whose innervation is responsible for initiating the gastric response to stress (Lewis *et al*., 2002). PVN^CRF^ projections were also investigated using the same methods as stated above. Following pHFD, there was a significant increase in CRF+ neurons in the posterior PVN compared to control rats (19.85±12.89 vs. 9.83±5.184 cells/section in pHFD and control rats, respectively, p=0.0158, t(30)=2.558, 95% C.I.=2.019 to 18.01; two-tailed unpaired t-test; Figure 4G). In contrast, there was no change in number PVN^CRF^ neurons in the anterior (27.00±8.367 vs. 18.88±17.22 cells/section in pHFD and control rats, respectively, p=0.4007, t(10)=0.8778, 95% C.I.=-12.50 to 28.75; two-tailed unpaired t-test; Figure 4C) or intermediate (50.25±13.85 vs. 53.36±21.04 cells/section in pHFD and control rats, respectively, p=0.6665, t(24)=0.4363, 95% C.I.=-17.81 to 11.59; two-tailed unpaired t-test; Figure 4E) PVN. Furthermore, there were no significant changes in numbers PVN^CRF^-DMV neurons in either the posterior (1.800±1.473 vs. 2.417±1.929 cells/section in pHFD and control rats, respectively, p=0.3155, t(30)=1.021, 95% C.I.=-1.850 to 0.6171; two-tailed unpaired t-test; Figure 4H), anterior (0.50±0.58 vs. 0.25±0.46 cells/section in pHFD and control rats, respectively, p=0.4332, t(10)=0.8165, 95% C.I.=-0.4322 to 0.9322; two-tailed unpaired t-test; Figure 4D) or intermediate (1.750±1.36 vs. 2.857±1.99 cells/section in pHFD and control rats, respectively, p=0.1172, t(24)=1.625, 95% C.I.=-2.513 to 0.299; two-tailed unpaired t-test; Figure 4F) PVN.

**FIGURE 4:**
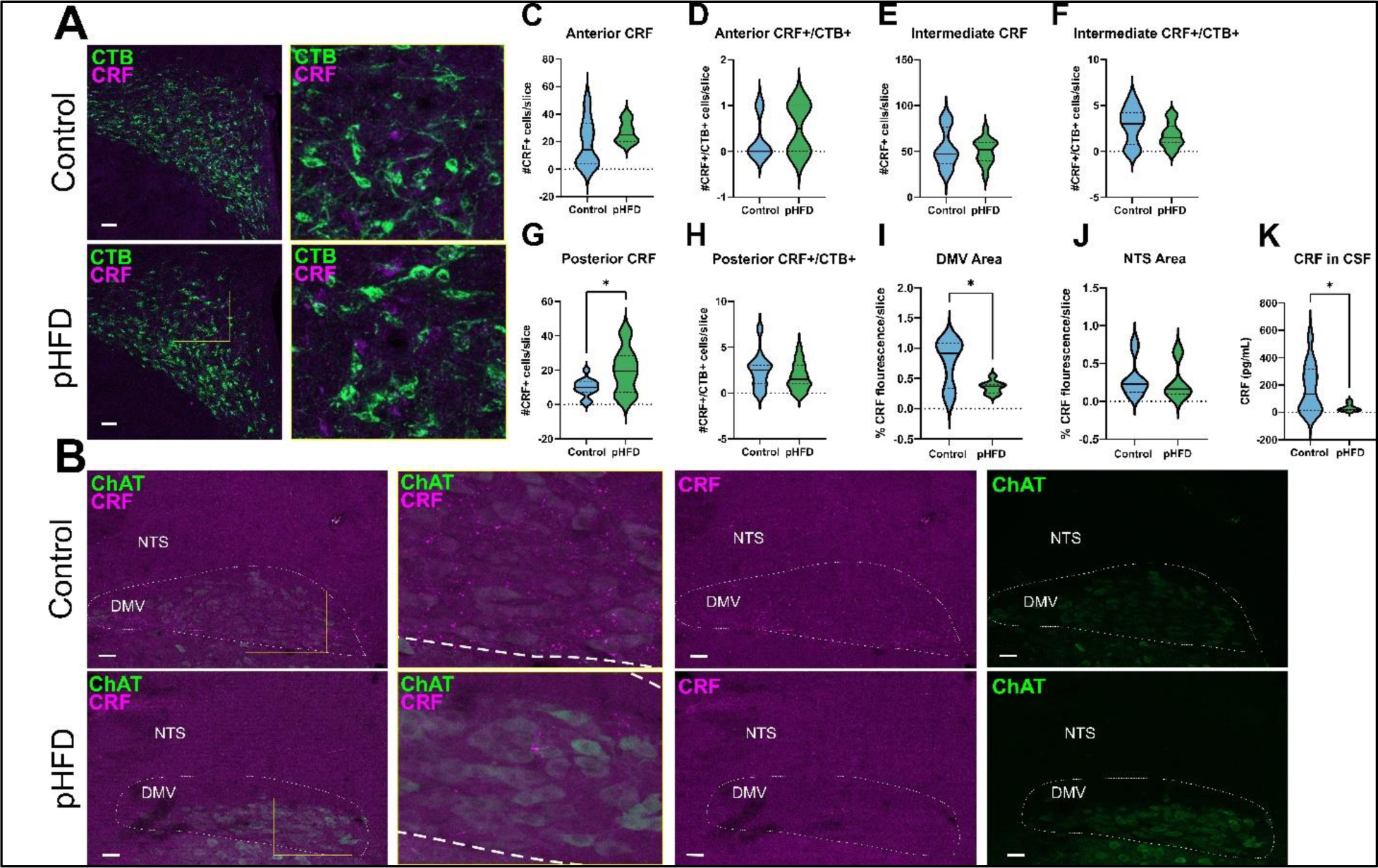
pHFD exposure increases PVN^CRF^-DMV neurons but reduces CRF-IR fibers in the DMV. **A:** Representative images of the posterior PVN with CTB+ neurons in green and CRF+ neurons in magenta from a control (top) and pHFD (bottom) rat illustrating the pHFD-induced increase in CRF+ neurons. Scale bar = 50 µm. **B:** Representative images of the brainstem containing the DMV (outlined; visualized via ChAT-IR to identify cholinergic neurons) and NTS in control (top) and pHFD (bottom) rats, illustrating the loss of CRF-IR fibers in the DMV following pHFD. Scale bar = 50µm. Quantification of CRF+ neurons (**C**) and CRF+/CTB+ neurons (**D**) in the anterior PVN show no significant difference between diet groups (N=4/diet). Similarly, quantification of CRF+ neurons (**E**) and CRF+/CTB+ neurons (**F**) in the intermediate PVN show no significant difference between diet groups. **G:** Graphical summary of the number of CRF+ neurons in the posterior PVN indicate a significant increase in pHFD compared to control rats (* p<0.05, two-tailed unpaired t-test). **H:** Graphical summary of the number of CRF and CTB co-localized neurons in the anterior PVN, showing no significant difference between pHFD and control rats. **I:** Graphical summary of % CRF fluorescence in the DMV show a significant reduction in pHFD rats (N=4) compared to controls (N=4) (* p<0.05, two-tailed unpaired t-test). **J:** Graphical summary of % CRF fluorescence in the NTS showing there is no difference between pHFD and control rats (N=4/diet). **K:** Graphical summary of CRF levels within CSF samples taken from pHFD (N=9) and control (N=13) rats demonstrating a significant decrease in CRF protein following pHFD. (* p<0.05, two-tailed unpaired t-test).

As above, immunohistochemistry was used to quantify CRF fibers at the intermediate level of the DVC (Bregma level −7.34) in both the DMV and NTS. pHFD exposure decreased CRF fiber intensity in the DMV (0.3575±0.1023% vs 0.7681±0.3851% in pHFD and control rats, respectively; p=0.0101, t(16)=2.918, 95% C.I.=-0.7089 to −0.1123, two-tailed unpaired t-test; Figure 4I) but not in the NTS (0.2615±0.2447% vs 0.2996±0.2468% in pHFD and control rats, respectively; p=0.7481, t(16)=0.3267, 95% C.I.=-0.2853 to 0.2091, two-tailed unpaired t-test; Figure 4J).

Lastly, CRF protein levels within 4^th^ ventricular CSF samples were assessed from both control (N=13) and pHFD (N=9). Compared to control rats, there was a significant reduction CSF CRF in pHFD rats (33.06±25.93 vs 178.6±177.0 pg/mL in pHFD and control rats, respectively; p=0.0246, t(20)=2.431, 95% C.I.=-270.5 to −20.67; via a two-tailed unpaired t-test, Figure 4K).

These results suggest that there is an overall increase in PVN^CRF^ neurons in rats exposed to a pHFD, although this does not translate to either the density of CRF fiber inputs into the DMV or to the CSF levels of CRF protein, although it should be noted that chronic stress exposure has been shown previously to decrease CRF levels within the CSF (Chappell *et al*., 1986).

### pHFD dysregulates gastric motility in a CRF-dependent manner

OXT and CRF have been shown to modulate gastric motility and tone in a stress-dependent manner (Stengel & Taché, 2010; Travagli & Anselmi, 2016). Specifically, DMV microinjection of OXT decreases corpus tone in control (unstressed) rats (Holmes *et al*., 2013) but this OXT-induced gastroinhibition is either attenuated or, in some instances, reversed following either acute stress or DMV application of CRF, likely due to short-term adaptive neuroplasticity to counteract the stress-induced decreases in gastric tone and motility (Travagli & Anselmi, 2016).

As shown previously, OXT induced a reduction in corpus tone in control rats (−134.2±79.06 mg; Figure 5C, D). In contrast, pHFD rats responded to OXT microinjection with a significant increase in gastric tone (+142.2±244.5 mg, p=0.0070, t(13)=3.201, 95% C.I.=89.84 to 462.9; two-tailed unpaired t-test; Figure 5C, D). There was no significant change in motility in response to OXT in either group (96.7±21.58 vs. 129.5±91.68 change from baseline motility, p=0.3138, t(13)=1.048, 95% C.I.=--34.79 to 100.3; two-tailed unpaired t-test; Figure 5E).

**FIGURE 5:**
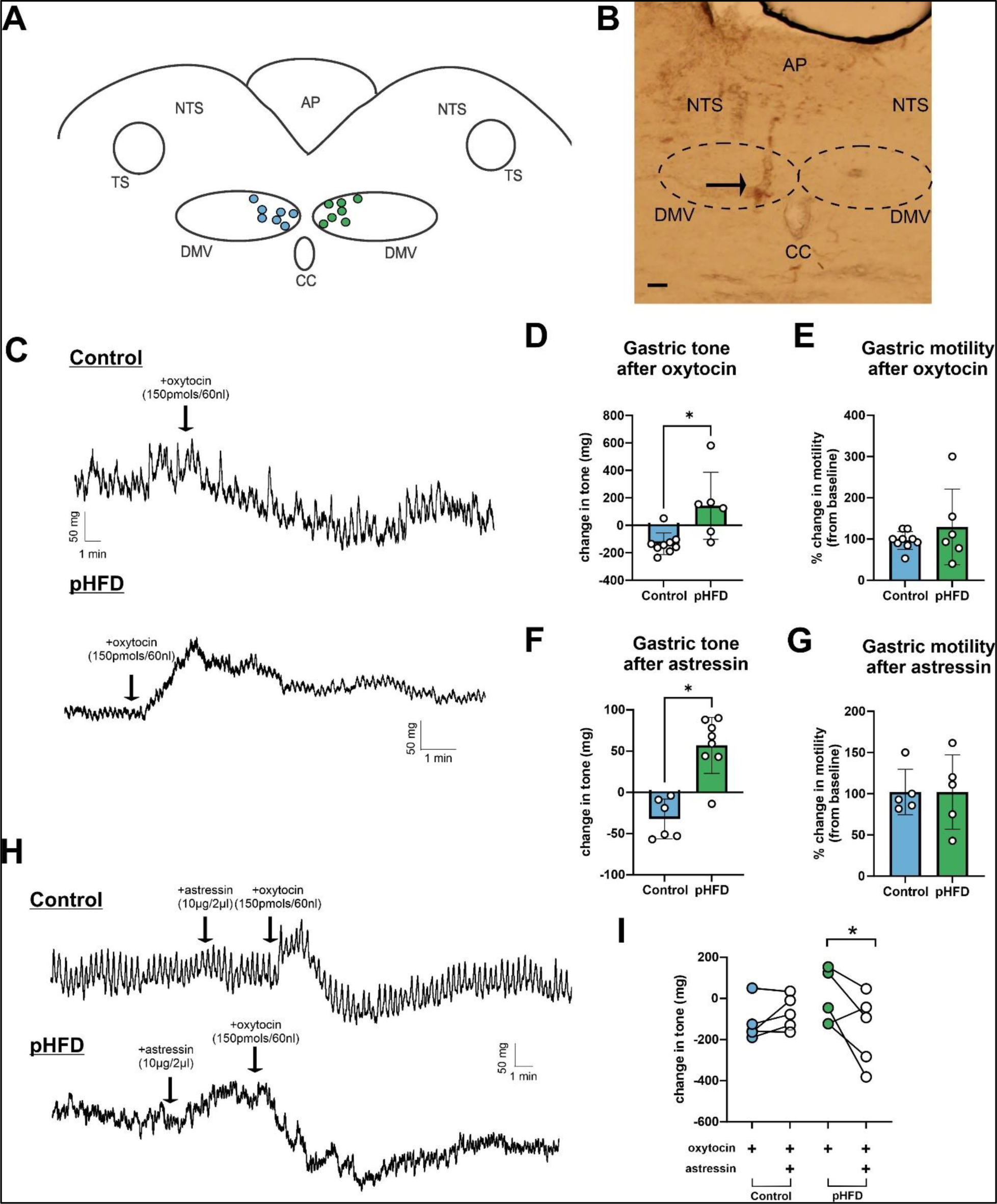
pHFD dysregulates gastric motility in a CRF-dependent manner. **A:** Schematic showing the microinjection sites of OXT in control (N=7, light blue circles, left) and pHFD (N=7, light green circles, right) animals for *in vivo* recordings of gastric corpus motility. All injections were made on the left side of the DMV but are separated out on the schematic for clarity. **B**: A representative image of a coronal section of the brainstem showing the microinjection pipette tract used to verify injection site (arrow). Scale bar = 50 µm. **C:** Representative traces of *in vivo* recordings of corpus gastric tone and motility in a control (top) and pHFD (rat) showing the larger basal motility in control rats as well as the inhibition in corpus tone following DMV OXT microinjection, but an increase in tone following DMV OXT microinjection in pHFD rats. **D:** Graphical summary of the significant difference in the OXT-induced change in corpus tone (in mg) in control (N=9) and pHFD (N=6) rats (two-tailed unpaired t-test * p<0.05). **E:** Graphical summary of the change in motility (as a % of the baseline motility) in control (N=9) and pHFD (N=6) rats showing no significant differences between diet groups. **F:** Graphical summary of the significant differences in corpus tone (in mg) in response to 4^th^ ventricular astressin application in control (N=6) and pHFD (N=8) (* p<0.05, two-tailed unpaired t-test). **G:** Graphical summary showing no change in motility in response to astressin application in control (N=5) and pHFD (N=5) rats. **H:** Representative *in vivo* recordings of corpus gastric tone and motility in a control (top) and pHFD (bottom) rat after 4^th^ ventricular application of astressin followed by a DMV OXT microinjection. Note that, in the control rat, astressin itself had no effect on tone, and did not alter the OXT-induced gastroinhibition. In contrast, in the pHFD rat, astressin itself increased corpus tone, and subsequent OXT microinjection led to a gastroinhibition. **I:** Graphical summary of the change in corpus tone in control and pHFD rats before and after astressin application (* p<0.05, two-tailed paired t-test).

Given the dysregulated response to OXT observed in pHFD rats, which is notably similar to that observed in control rats after stress (Jiang & Travagli, 2020), we investigated whether the observed differences were caused by CRF. To investigate this, the response to OXT was assessed before and after 4^th^ ventricular application of the CRFR1 antagonist, astressin. Astressin itself had no effect on gastric motility or tone in control rats (102.1±45.22 % change from baseline motility and 32.2±24.12 mg change from baseline tone, respectively, Figure 5F, G) and did not alter the gastroinhibitory response to OXT (−117.6±96.51 mg before and −69.4±82.35 mg after astressin treatment, p=0.1849, t(4)=1.600, 95% C.I.=-35.46 to 131.9; two-tailed paired t-test, Figure 5H, I) confirming that, as shown previously (Lewis *et al*., 2002), CRF receptors are not tonically active in control rats. In contrast, in pHFD rats, astressin increased corpus tone (+57.0±33.85 mg change from baseline; p=0.0001, t(12)=5.470. 95% C.I.=53.65 to 124.7; two-tailed unpaired t-test; Figure 5F) without affecting motility (102.1±27.68 % change from baseline, p=0.9992, t(8)=0.0.0011, 95% C.I.=--54.65 to 54.70; two-tailed unpaired t-test; Figure 5G). Furthermore, re-application of OXT after astressin pre-treatment led to a significant decrease in tone in pHFD rats (+142.2±244.5 mg before and −75. 7±271.67 mg after astressin treatment, p=0.0015, t(5)=6.249, 95% C.I.=-307.4 to −128.2; two-tailed paired t-test, Figure 5H, I). Indeed, the OXT-induced gastroinhibition observed in pHFD rats following astressin pretreatment was not different from that observed in control rats (−75. 7±271.4 mg change from baseline vs. −69.4±82.35 mg change from baseline in pHFD and control rats, respectively, p=0.9617, t(9)=0.04938, 95% C.I.=--293.3 to 280.8; two-tailed unpaired t-test; Figure 5I) suggesting that, in pHFD rats, antagonizing tonically active CRF receptors normalizes vagally-dependent responses to OXT.

Taken together, these results suggest that the tonic activation of CRF receptors in pHFD rats is responsible for the dysregulated gastric responses to OXT, further supporting the hypothesis that pHFD exposure induces maladaptive neuroplasticity within brainstem vagal circuits regulating gastric functions.

### pHFD uncovers OXT-mediated inhibition of GABAergic synaptic currents in a CRF-dependent manner

Previous studies have shown that OXT has no effect on inhibitory GABAergic synaptic transmission to gastric-projecting DMV neurons unless rats are exposed to acute stress or CRF (Browning *et al*., 2014). Specifically, following acute stress (or CRF exposure), OXT decreases GABAergic NTS-DMV transmission, disinhibiting vagal efferent drive to the stomach and increasing gastric tone and motility (Holmes *et al*., 2013; Browning *et al*., 2014; Jiang & Travagli, 2020). To determine if the dysregulated gastric responses observed in pHFD rats are due to the uncovering of OXT-mediated effects, whole cell patch clamp recordings were made from DMV neurons in control (N=16) and pHFD (N=20) rats to assess changes in GABAergic synaptic transmission.

As shown previously, control DMV neurons showed no change in mIPSC properties, either in mIPSC frequency (99.3±14.72 % of baseline; p=0.8901, t(8)=0.1427, 95% C.I.=-12.02 to 10.62; two-tailed one sample t-test (µ=100 %), N = 9 neurons from 5 rats; Figure 6A, B) or charge transfer (105.1±30.92 % of baseline; p=0.6370, t(8)=0.4904, 95% C.I.=-18.71 to 28.82; two-tailed one sample t-test (µ=100%), N = 9 neurons from 5 rats; Figure 6C) (see Table 1 for details). Interestingly, control DMV neurons showed a significant change in mIPSC amplitude following OXT application (91.1±10.29 % of baseline, p=0.0323, t(8)=2.585, 95% C.I.=-16.78 to −0.9590; via two-tailed one sample t-test (µ=100 %), N = 9 neurons from 5 rats; Figure 6D), however the magnitude of the change (−8.872 %) would suggest this is unlikely to be of physiological significance and previous studies defined ‘responding neurons’ as those in which an effect >25% was observed.

**FIGURE 6:**
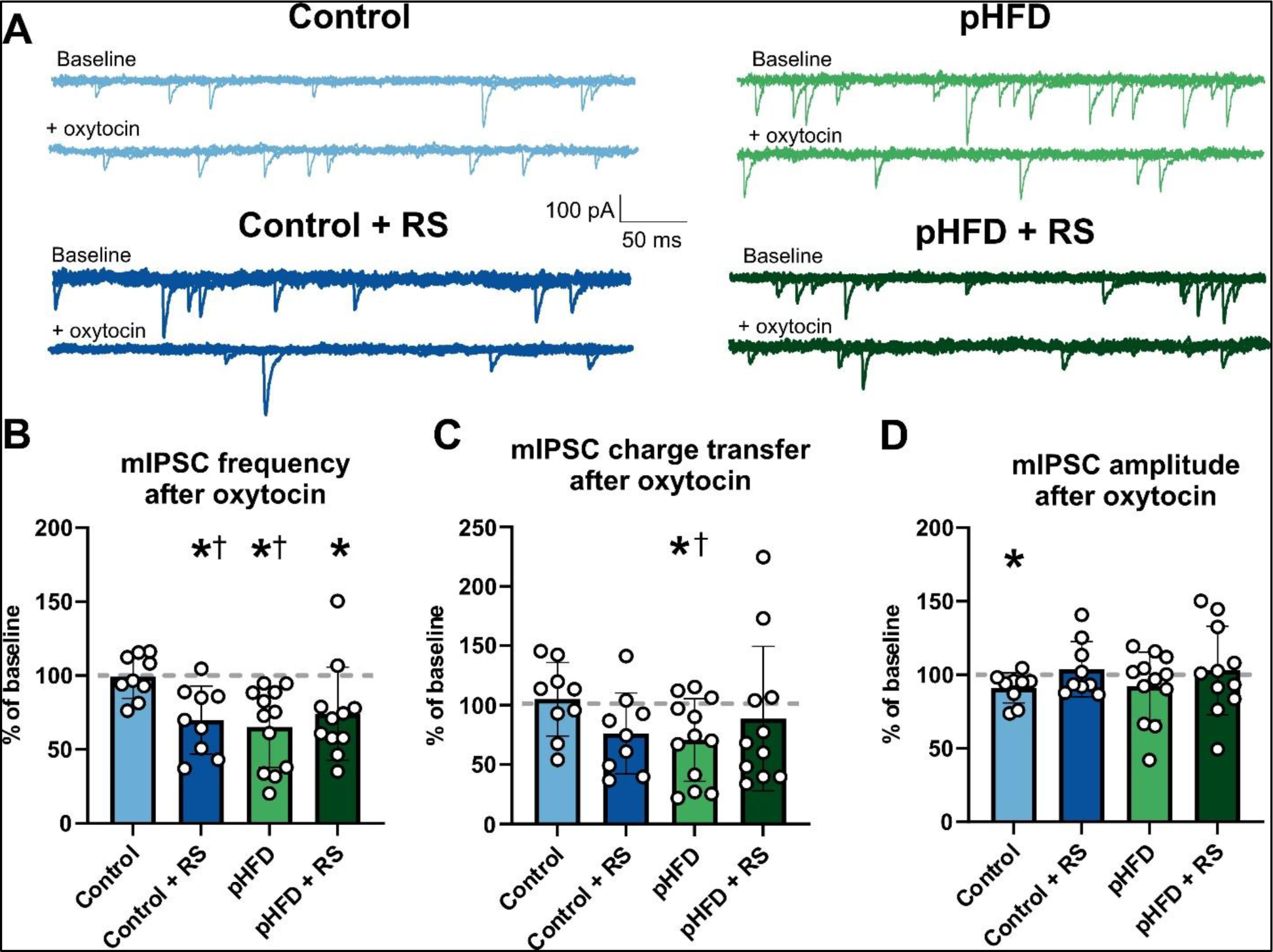
pHFD uncovers OXT-mediated inhibition of GABAergic synaptic currents. **A:** Representative overlapping traces of miniature inhibitory post synaptic currents (mIPSCs) recorded from control (light blue), pHFD (light green), control + RS (dark blue), and pHFD + RS (dark green) DMV neurons before (top trace) and after (bottom trace) application of OXT. Baseline conditions were recorded in neurons voltage clamped at −50mV in the presence of TTX and kynurenic acid. Note that in control neurons, OXT has no effect on mIPSC, but decreases mIPSC frequency following RS. In contrast, OXT decreases mIPSC frequency in unstressed pHFD neurons and acute stress has no additional effects. **B:** Graphical summary of the OXT-induced change in mIPSC frequency in control (N=9), control + RS (N=9), pHFD (N=12), and pHFD + RS (N=11) neurons, demonstrating the inhibitory effects of OXT after RS in control neurons, and following pHFD as well as pHFD + RS (* p<0.05 two-tailed one sample t-test against baseline; † p<0.05 two-tailed unpaired t-test compared to the naive control group). **C:** Graphical summary of the change in charge transfer of mIPSCs after OXT application illustrating that pHFD exposure-induced inhibition (* p<0.05 two-tailed one sample t-test against baseline; † p<0.05 two-tailed unpaired t-test compared to the naïve control group). **D:** Graphical summary of the change in mIPSC amplitude after OXT application (* p<0.05 two-tailed one sample t-test vs baseline).

**TABLE 1:** effects of Oxytocin on mIPSC parameters.

| Group | Baseline | + Oxytocin (100 nM) |
| --- | --- | --- |
| **mIPSC frequency** |  |  |
| Control | 1.469 ± 1.108 Hz | 1.530 ± 1.317 Hz |
| Control + RS | 3.320 ± 3.157 Hz | 2.187 ± 2.430 Hz |
| pHFD | 2.153 ± 1.059 Hz | 1.408 ± 0.804 Hz |
| pHFD + RS | 2.257 ± 1.623 Hz | 1.798 ± 1.681 Hz |
| **mIPSC amplitude** |  |  |
| Control | 65.448 ± 14.891 pA | 59.022 ± 12.054 pA |
| Control + RS | 51.870 ± 17.783 pA | 53.366 ± 17.908 pA |
| pHFD | 80.210 ± 19.317 pA | 71.866 ± 18.949 pA |
| pHFD + RS | 61.187 ± 30.144 pA | 61.908 ± 38.041 pA |
| **mIPSC charge transfer** |  |  |
| Control | 454.208 ± 421.153 pAms.s<sup>-1</sup> | 508.074 ± 617.797 pAms.s<sup>-1</sup> |
| Control + RS | 814.720 ± 989.933 pAms.s<sup>-1</sup> | 456.454 ± 451.126 pAms.s<sup>-1</sup> |
| pHFD | 591.747 ± 378.469 pAms.s<sup>-1</sup> | 378.469 ± 232.087 pAms.s<sup>-1</sup> |
| pHFD + RS | 917.548 ± 946.005 pAms.s<sup>-1</sup> | 982.110 ± 1533.16 pAms.s<sup>-1</sup> |

As shown previously, acute stress uncovered the ability of OXT to inhibit GABAergic synaptic transmission in control rats, decreasing mIPSC frequency (69.9±23.03 % of baseline; p=0.0044, t(8)=3.916, 95% C.I.=-47.76 to −12.36; two-tailed one sample t-test (µ=100 %), N = 9 neurons from 4 rats; Figure 6A, B) without affecting mIPSC charge transfer (76.3±34.05 % of baseline; p=0.0700, t(8)=2.091, 95% C.I.=-49.91 to 2.445; two-tailed one sample t-test (µ=100 %), Figure 6C) or mIPSC amplitude (103.8±18.79 % of baseline; p=0.5596, t(8)=0.6086, 95% C.I.=-10.63 to 18.26; two-tailed one sample t-test (µ=100 %), N = 9 neurons from 4 rats; Figure 6D). Restraint stress significantly changed the effects of OXT effects on frequency in control diet conditions (69.94±23.03 % vs 99.30±14.72 % of baseline; p=0.0053, t(16)=3.223. 95% C.I.=-48.68 to −10.05; via two-tailed unpaired t-test, N = 9 neurons from 4 rats; Figure 6B).

In naive (i.e. unstressed) pHFD neurons, however, OXT decreased mIPSC frequency (65.2±27.30 % of baseline; p=0.0010, t(11)=4.418, 95% C.I.=-52.17 to −17.47; two-tailed one sample t-test (µ=100 %), N = 12 neurons from 7 rats; Figure 6A, B) and charge transfer (70.7±34.76 % of baseline; p=0.0138, t(11)=2.924, 95% C.I.=-51.43 to −7.254; two-tailed one sample t-test (µ=100), Figure 6C), without affecting mIPSC amplitude (92.2±23.22 % of baseline; p=0.2718, t(11)=1.157, 95% C.I.=-22.51 to 6.998; two-tailed one sample t-test (µ=100), N = 12 neurons from 7 rats; Figure 6D; see Table 1 for details). The OXT-induced reduction in mIPSC frequency and charge transfer was significantly different than that observed in naive controls (mIPSC frequency: 65.2±27.30 % vs 99.3±14.72 % of baseline; p=0.0031, t(19)=3.384, 95% C.I.=-55.22 to −13.02; two-tailed unpaired t-test; charge transfer: 70.7±34.76 % vs 105.1±30.92 % of baseline; p=0.0298, t(19)=2.350. 95% C.I.=-65.03 to −3.755; two-tailed unpaired t-test, Figure 6C). Following stress, OXT still induced a significant decrease in GABAergic transmission in pHFD rats (mIPSC frequency: 74.3±31.51 % of baseline; p=0.0220, t(10)=2.708, 95% C.I.=-46.90 to −4.563; two-tailed one sample t-test (µ=100 %), N = 11 neurons from 5 rats; Figure 6B), but this effect was not different from that induced in naive pHFD neurons (74.3±31.51 % vs 65.2±27.30 % of baseline; p=0.4668, t(21)=0.7412, 95% C.I.=-16.42 to 34.59; two-tailed unpaired t-test; Figure 6B)

These results suggest that, like acute stress, pHFD uncovers the ability of OXT to inhibit GABAergic transmission at the NTS-DMV synapse and, further, that, contrary to what observed in control rats, stress has no additional effects in pHFD rats.

To determine whether the ability of OXT to decrease GABAergic synaptic transmission following pHFD was due to tonic activation of CRF receptors, the effects of astressin on mIPSCs and its ability to prevent or attenuate the OXT-mediated inhibition was assessed. In naïve (i.e. unstressed) control DMV neurons, astressin application led to a statistically (although likely not physiologically) significant decrease in mIPSC frequency (83.3±5.585 % of baseline; p=0.0002, t(7)=7.911, 95% C.I.=-21.87 to −11.53; two-tailed one sample t-test (µ=100), N = 7 neurons from 3 rats; Figure 7B) and a significant decrease in mIPSC charge transfer (69.1±16.02 % of baseline; p=0.0052, t(5)=4.724, 95% C.I.=-47.72 to −14.09; two-tailed one sample t-test (µ=100 %), N = 7 neurons from 3 rats; Figure 7C) with no effect on mIPSC amplitude (94.9±11.28 % of baseline; p=0.2801, t(7)=1.187, 95% C.I.=-15.49 to 5.372; two-tailed one sample t-test (µ=100 %), N = 7 neurons from 3 rats; Figure 7D, see Table 2 for details).

**FIGURE 7:**
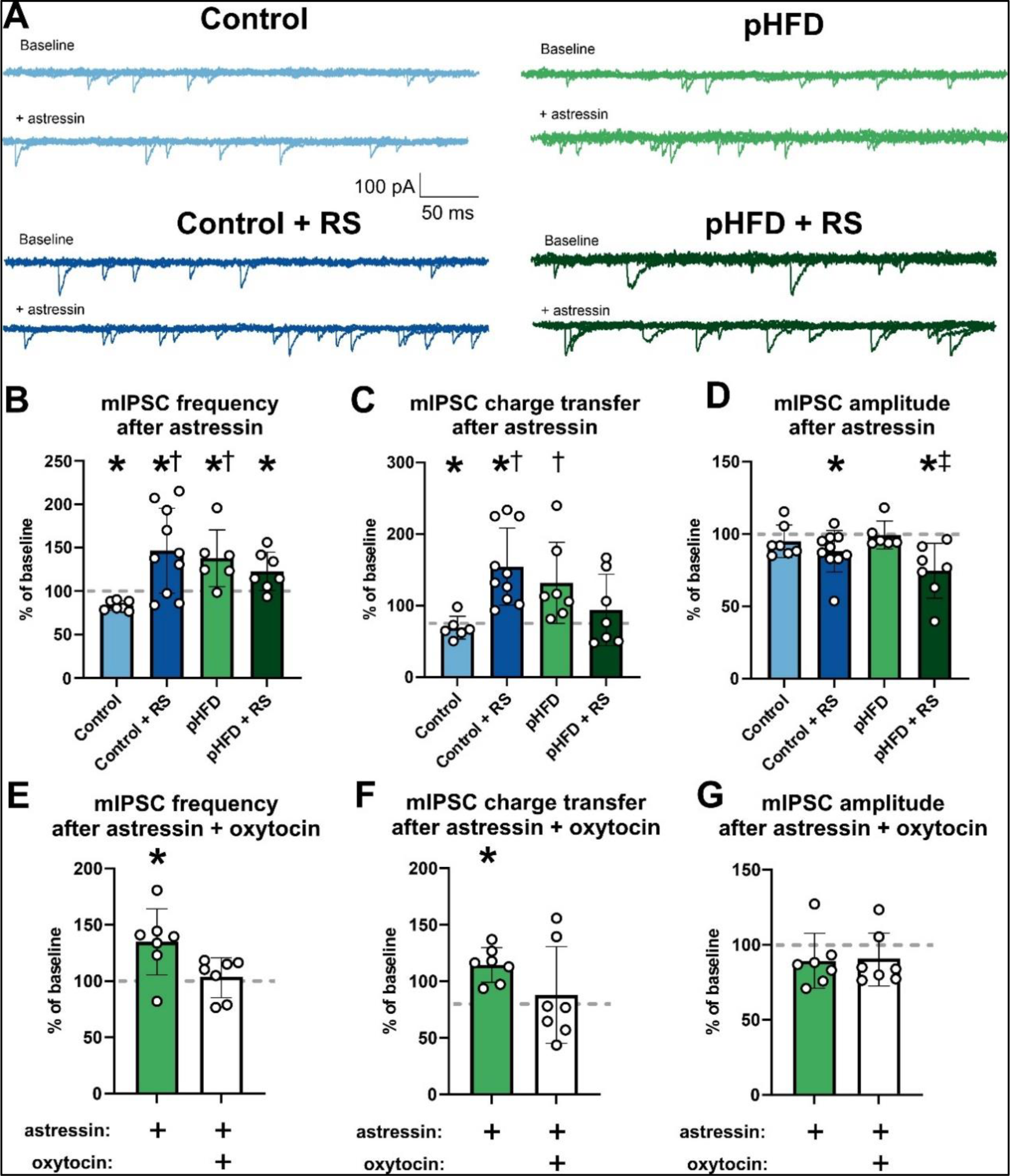
pHFD tonically activates CRF1 receptors at NTS-DMV synapses. **A:** Representative traces of miniature inhibitory post synaptic currents (mIPSCs) recorded from control (light blue), pHFD (light green), control + RS (dark blue), and pHFD + RS (dark green) DMV neurons before (top trace) and after (bottom trace) application of astressin. mIPSCs were recorded in DMV neurons voltage clamped at −50mV in the presence TTX and kynurenic acid. Note that in control rats astressin has decreased mIPSCs frequency under naive conditions, but increases mIPSC frequency following acute stress. In contrast, astressin application increased mIPSC frequency in pHFD rats with no additional effect after the RS. Graphical summary of the astressin-induced changes in mIPSC frequency (**B**), charge transfer (**C**) and amplitude (**D**), in control (N=7), control + RS (N=10), pHFD (N=6), and pHFD + RS (N=7) neurons (* p<0.05 using a two-tailed one sample t-test vs baseline; † p<0.05 using a two-tailed unpaired t-test compared to the unstressed control group). Graphical summary of results assessing whether astressin attenuated the ability of OXT to inhibit mIPSC frequency (**E**), charge transfer (**F**) and amplitude (**G**). p<0.05 using a two-tailed one sample t-test vs baseline).

**TABLE 2:** Effects of Astressin on mIPSC parameters.

| Group | Baseline | + Astressin (1 μM) |
| --- | --- | --- |
| **mIPSC frequency** |  |  |
| Control | 1.791 ± 0.921 Hz | 1.455 ± 0.667 Hz |
| Control + RS | 2.025 ± 1.066 Hz | 2.927 ± 1.638 Hz |
| pHFD | 1.042 ± 0.437 Hz | 1.309 ± 0.470 Hz |
| pHFD + RS | 1.432 ± 0.707 Hz | 1.675 ± 0.700 Hz |
| **mIPSC amplitude** |  |  |
| Control | 67.824 ± 17.946 pA | 64.048 ± 15.730 pA |
| Control + RS | 67.456 ± 16.480 pA | 58.963 ± 15.687 pA |
| pHFD | 56.395 ± 21.119 pA | 55.106 ± 21.736 pA |
| pHFD + RS | 54.499 ± 24.306 pA | 39.955 ± 17.735 pA |
| **mIPSC charge transfer** |  |  |
| Control | 975.225 ± 899.930 pAms.s<sup>-1</sup> | 676.028 ± 562.638 pAms.s<sup>-1</sup> |
| Control + RS | 611.678 ± 318.242 pAms.s<sup>-1</sup> | 967.957 ± 695.137 pAms.s<sup>-1</sup> |
| pHFD | 295.478 ± 141.286 pAms.s<sup>-1</sup> | 349.979 ± 130.904 pAms.s<sup>-1</sup> |
| pHFD + RS | 599.292 ± 421.968 pAms.s<sup>-1</sup> | 435.281 ± 237.549 pAms.s<sup>-1</sup> |

Following acute stress, however, in control neurons astressin caused a significant increase in mIPSC frequency (146.5±48.81 % of baseline; p=0.0146, t(9)=3.014, 95% C.I.=11.60 to 81.43; two-tailed one sample t-test (µ=100 %), N = 7 neurons from 4 rats; Figure 7B) and charge transfer (154.5±53.95 % of baseline; p=0.0109, t(9)=3.194, 95% C.I.=15.90 to 93.09; two-tailed one sample t-test (µ=100 %), N = 7 neurons from 4 rats; Figure 7C), with a statistically significant decrease in mIPSC amplitude (88.04±14.37 % of baseline; p=0.0273, t(9)=2.632, 95% C.I.=-22.24 to −1.679; two-tailed one sample t-test (µ=100 %), N = 7 neurons from 4 rats; Figure 7D) in control DMV neurons. These astressin-induced effects were significantly different in stressed vs naive control neurons (mIPSC frequency: 146.5±48.81 % vs 83.3±5.585 % of baseline; p=0.0041, t(15)=3.378, 95% C.I.=23.33 to 103.1; two-tailed unpaired t-test; and charge transfer: 154.5±53.95 % vs 69.1±16.02 % of baseline; p=0.0022, t(14)=3.733, 95% C.I.=36.33 to 134.5; two-tailed unpaired t-test).

In contrast, in naive pHFD neurons, astressin itself increased mIPSC frequency (132.4±33.06 % of baseline; p=0.0409, t(6)=2.596, 95% C.I.=1.861 to 63.01; two-tailed one sample t-test (µ=100 %), N = 10 neurons from 5 rats; Figure 7B) without affecting charge transfer (131.7±56.65 % of baseline; p=0.1895, t(6)=1.480, 95% C.I.=-20.71 to 84.08; two-tailed one sample t-test (µ=100 %), N = 10 neurons from 5 rats; Figure 7C) or amplitude (99.3±9.649 % of baseline; p=0.8662, t(5)=0.1774, 95% C.I.=-10.82 to 9.427; two-tailed one sample t-test (µ=100 %), N = 10 neurons from 5 rats; Figure 7D). The astressin-mediated effects on GABAergic transmission in naive pHFD neurons were significantly different from those observed in naive control neurons (mIPSC frequency: 132.4±33.06 % vs 83.3±5.585 % of baseline; p=0.0022, t(12)=3.877, 95% C.I.=21.52 to 76.75; two-tailed unpaired t-test, Figure 7B; charge transfer: 131.7±56.65% vs 69.1±16.02 % of baseline; p=0.0245, t(11)=2.603, 95% C.I.=9.672 to 115.5; two-tailed unpaired t-test, Figure 7C).

Further, following acute stress, pHFD neurons still responded to astressin with an increase in mIPSC frequency (122.8±22.04 % of baseline; p=0.0338, t(6)=2.739, 95% C.I.=2.430 to 43.20; two-tailed one sample t-test (µ=100 %), N = 7 neurons from 4 rats; Figure 7B) with no change to mIPSC charge transfer (93.9±50.03 % of baseline; p=0.7591, t(6)=0.3209, 95% C.I.=-52.34 to 40.20; two-tailed one sample t-test (µ=100%), N = 7 neurons from 4 rats; Figure 7C), and an astressin-mediated decrease in mIPSC amplitude (74.6±19.02 % of baseline; p=0.0123, t(6)=3.534, 95% C.I.=-43.00 to −7.817; two-tailed one sample t-test (µ=100 %), N = 7 neurons from 4 rats; Figure 7D), but these responses were not different from those observed in naive pHFD neurons (mIPSC frequency: 122.8±22.04 % vs 132.4±33.06 % of baseline; p=0.5337, t(12)=0.6408, 95% C.I.=-42.35 to 23.10; two-tailed unpaired t-test, Figure 7B; charge transfer: 93.9±50.03 % vs 131.7±56.65 % of baseline, p=0.2110, t(12)=1.321, 95% C.I.=-99.99 to 24.49; two-tailed unpaired t-test, Figure 7C). It should be noted, however, that the astressin-induced changes in mIPSC amplitude in pHFD neurons were significantly different following stress (74.6±19.02 % vs 99.3±9.649 % of baseline; p=0.0153, t(12)=2.869, 95% C.I.=-43.67 to −5.753; two-tailed unpaired t-test, Figure 7D).

To determine if the OXT-mediated decrease in mIPSCs observed following pHFD exposure was driven by tonic activation of CRF1 receptors at the level of the NTS-DMV synapse, the effects of OXT were assessed following astressin application. As before, astressin increased mIPSC frequency (135.0±29.35 % of baseline; p=0.0197, t(6)=3.154, 95% C.I.=7.840 to 62.12; via two-tailed one sample t-test (µ=100 %), N = 7 neurons from 4 rats; Figure 7E) and charge transfer (114.6±15.31 % of baseline; p=0.0448, t(6)=2.527, 95% C.I.=0.4646 to 28.77; via two-tailed one sample t-test (µ=100 %), N = 7 neurons from 4 rats; Figure 7F, see Table 3 for details). Subsequent application of OXT in the presence of astressin, however, no longer induced a significant change in mIPSC frequency (103.0±17.78 % of baseline; p=0.6710, t(6)=0.4463, 95% C.I.=-13.45 to 19.45; via two-tailed one sample t-test (µ=100 %), Figure 7E) or charge transfer (87.99±42.72 % of baseline; p=0.4850, t(6)=0.7439, 95% C.I.=-51.53 to 27.50; via two-tailed one sample t-test (µ=100 %), N = 7 neurons from 4 rats; Figure 7F), indicating that the OXT-mediated decrease in GABAergic transmission was driven by tonic CRF receptor activation.

**TABLE 3:**
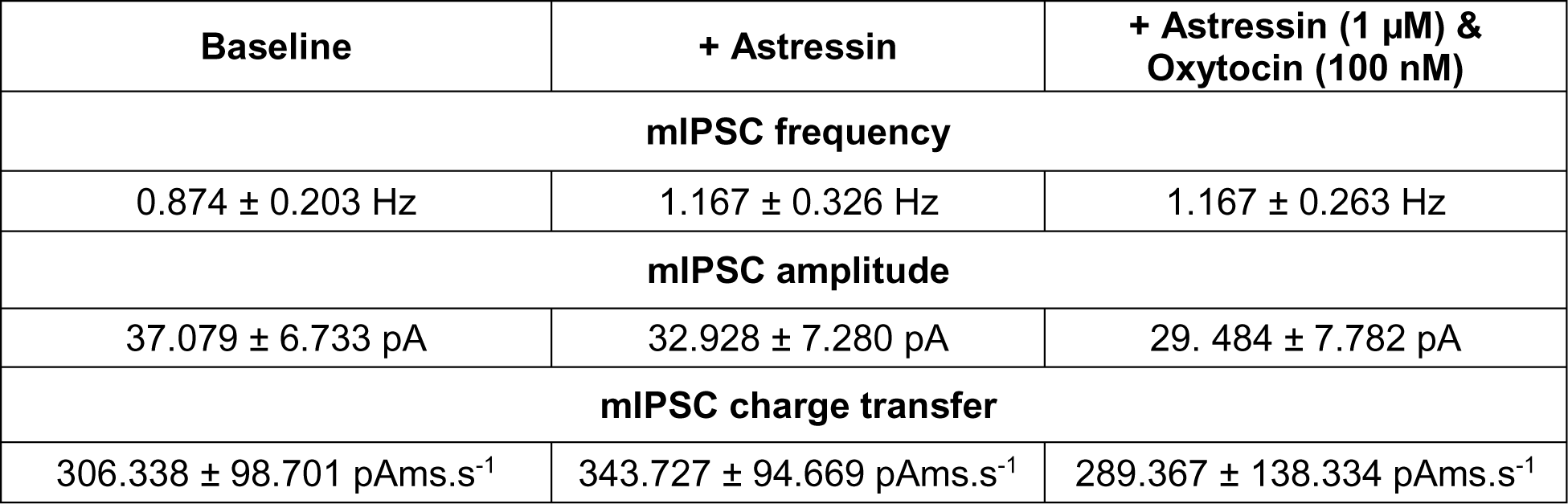
Effects of Astressin to modulate oxytocin effects on mIPSC parameters in pHFD DMV neurons.

Taken together, these results demonstrate that, similar to acute stress, pHFD induces tonic release of CRF and activation of CRF1 receptors at the NTS-DMV synapse.

## DISCUSSION

The results from the present study support the hypothesis that pHFD alters descending PVN-DMV inputs and dysregulates vagal brain-gut responses to stress.

In particular, our novel data suggest that pHFD exposure (i) delayed gastric emptying and prevented rats from mounting the appropriate gastroinhibitory response to acute stress, (ii) altered PVN^CRF^ and PVN^OXT^ projections to the DMV which was responsible for (iii) tonic activation of DVC CRF1 receptors and (iv) dysregulated gastric motility and vagally-mediated responses to OXT.

Taken together, these results suggest that pHFD alters the development of PVN-DMV neurocircuitry such that pHFD rats exhibit a “stressed” vagal phenotype which may contribute to a lack of resiliency and dysregulated gastric responses to stress.

An acute stressor, defined as an external event perceived to threaten homeostasis, leads to activation of the hypothalamic-pituitary-adrenal (HPA) axis (Chrousos, 2009). The HPA axis contains both neural and endocrine components that must coordinate their responses to result in glucocorticoid production and release into circulation. Beyond this defined circuit, both the neural (the autonomic nervous system) and endocrine (circulating hormones) components act on various effector organs to coordinate the organism’s adaptive response to stress. This includes activation of neural networks that elevate cardiovascular function and blood glucose production, increased attention/focus, in addition to inactivating non-essential functions such as digestion and reproduction (Chrousos & Gold, 1992; Chrousos, 2009). Once the threat, (real or perceived) is resolved, this neuroendocrine system attenuates these stress-induced adaptations to reestablish homeostasis. Specifically, elevated levels of glucocorticoids act as one source of negative feedback, shutting down HPA axis activation (Herman *et al*., 2016; Gjerstad *et al*., 2018). A rise in CORT is expected after an acute stress exposure, however in the current study serum samples were taken at the end of the 2-hour restraint stress, so in control rats their CORT levels have returned to baseline whereas pHFD rats still have elevated CORT levels. This suggest that, following pHFD, the endocrine component of the HPA circuit appears to be dysfunctional, as circulating corticosterone levels remain elevated following the restraint stress, potentially suggesting a loss of negative feedback mechanisms or an increased HPA circuit drive. Of note, results of the current study also demonstrated that basal blood glucose levels were elevated in following pHFD, and the stress-induced increase in glycemic level was also exaggerated. Certainly, diet-induced obesity is associated with an increase in blood glucose level as insulin insensitivity develops (Blázquez & Quijada, 1968; Lichtenstein & Schwab, 2000). While, notably, the pHFD rats used in the current study were not obese, it is nevertheless likely they were insulin-intolerant, if not yet insulin resistant (Czech, 2017; Erion & Corkey, 2017; Kislal *et al*., 2020).

In contrast, as evidenced by the delay in gastric emptying even at baseline and in the absence of stress, pHFD uncouples the parasympathetic GI stress response from the neuroendocrine stress response. Regarding many of these ‘stress’ markers, including *in vitro* DMV neuronal responses, *in vivo* gastric emptying rates, and *in vivo* gastric motility, the current study suggests that pHFD rats appear to already be “stressed”.

Stress affects gastric functions in a vagally-dependent manner, specifically through CRF and OXT signaling from PVN parvocellular and magnocellular neurons, respectively, that innervate the DVC. Both CRF and OXT alter vagal efferent outflow directly (via actions to modulate DMV neurons; (Raggenbass *et al*., 1987; Dreifuss *et al*., 1992; Raggenbass & Dreifuss, 1992; Lewis *et al*., 2002)) as well as indirectly (via actions to modulate NTS-DMV synaptic inputs; (van den Pol, 1982; Wang *et al*., 1996; Holmes *et al*., 2013; Browning *et al*., 2014)). Exogenous application of CRF to the brainstem (or exposure to an acute stressor) delays gastric emptying (Martínez *et al*., 1997) and inhibits gastric tone (Lewis *et al*., 2002), whereas OXT acts to counteract these changes to gastric physiology (Holmes *et al*., 2013; Browning *et al*., 2014). Such gastroinhibitory responses to acute stress are protective at the level of the stomach (Browning & Travagli, 2019); equally important, however, is the ability to attenuate this inhibitory response when exposed to chronic stress (i.e., stress adaptation or resiliency). Previous studies have shown that Sprague Dawley rats adapt to chronic homotypic stress by upregulating PVN^OXT^-DMV projections, allowing normalization of gastric emptying rates and gastric tone (Jiang *et al*., 2018). The current study has shown that pHFD decreases PVN^OXT^-DMV projections; it remains to be determined whether this loss of OXT input plays a role in apparent inability to adapt to chronic stress, which manifests as increased anxiety and reduced stress resiliency in this population (Rivera *et al*., 2015; DeCapo *et al*., 2019). Of interest, acute stress elevates blood pressure via CRF-dependent projections to the NTS and subsequent alterations to sympathetic cardiac outflow (Wang *et al*., 2019) although it remains to be determined whether cardiovascular autonomic outputs are similarly uncoupled following pHFD. Additionally, while the current study has suggested that pHFD disrupts hypothalamic-brainstem stress neurocircuits that impact GI functions, investigation of limbic regions (the central amygdala and anterior cingulate cortex) and higher order regions involved in executive function (prefrontal cortex) would be required to assess the role of descending inputs to the hypothalamus as additional potential drivers of this neural dysfunction, given their roles in anxiety (Krieger *et al*., 2022) and pain perception (Silverman *et al*., 1997). These are also sources of extrahypothalamic CRF production (Merchenthaler *et al*., 1982; Bhatia & Tandon, 2005) and could explain the dissonance between PVN^CRF^ levels, CRF fiber density in the DMV, and CRF concentration in the CSF identified in the current study.

pHFD exposure has been shown previously to decrease DMV neuronal excitability and function, with notable effects to increase GABAergic signaling. Specifically, pHFD prevents the maturation of GABA_A_ receptor subunit phenotype, leading to slower GABA currents (Clyburn *et al*., 2019). Together with alterations in the DMV neuronal morphology (larger neurons, increased dendritic arborization) and membrane biophysical properties (lower input resistance, larger action potential afterhyperpolarization amplitude, reduced depolarization induced firing frequency) (Bhagat *et al*., 2015), this would exacerbate the pHFD induced decrease in neuronal excitability and vagal efferent output. The current study extends these phenotypic changes to include pHFD-induced alterations in the responsiveness of DMV neurons to neuroendocrine hormones, which appear, at least in part, due to the tonic activation of CRF1 receptors. Several studies have shown that OXT alters NTS-DMV GABAergic transmission only after stress or CRF exposure (Browning *et al*., 2014; Jiang *et al*., 2018); the ‘state of activation’ of inhibitory NTS-DMV synapses is low due to the tonic inhibition of adenylate cyclase which decreases cAMP-PKA signaling. Stress, or CRF, activates adenylate cyclase, increases cAMP-PKA signaling and uncovers the presynaptic inhibitory effects of OXT, the results of which is an engagement of vagal efferent pathways that overcome the gastroinhibitory response to stress (Browning *et al*., 2014). In contrast, pHFD uncovers the ability of OXT to inhibit GABAergic synaptic currents prior to stress. Indeed, this presynaptic inhibitory effect of OXT is normalized following CRF1 receptor antagonism suggesting this mechanism is already engaged in pHFD rats. Future studies will be required to determine the balance of excitatory and inhibitory vagal efferent pathways that modulate gastric functions in pHFD rats, as well as their potential to compensate for central alterations at the NTS-DMV synapse.

The current study suggests that DVC neurocircuits are primed by pHFD to respond to OXT, but the reduction in PVN^OXT^-DMV inputs prevents this response. Given that chemogenetic activation of oxytocin prevents stress-induced gastroinhibition (Jiang & Travagli, 2020), and stress resiliency and adaptation is associated with an upregulation of these inputs (Jiang *et al*., 2018), it will be important for future studies to investigate whether pHFD rats are capable of upregulating PVN^OXT^-DMV inputs and whether this would alleviate the basal gastric dysregulation. Of interest, nasal OXT application reduces social and separation anxiety (Buchheim *et al*., 2009; Alvares *et al*., 2010; de Oliveira *et al*., 2012; Clark-Elford *et al*., 2014) and improves social behavior in children with autism spectrum disorder (Huang *et al*., 2021); both of these disorders are associated with GI dysfunction (Holtmann *et al*., 2017; Azhari *et al*., 2019). Similarly, it will be important to determine the precise period during which diet exposure is critical for proper development; vagal circuits begin to develop at embryonic day 13 (Rinaman & Levitt, 1993), but continue to refine into the second postnatal week (Rao *et al*., 1997; Vincent & Tell, 1997), while descending hypothalamic-brainstem neurocircuits are not mature until weaning (Rinaman, 2003, 2006).

Studies looking at the offspring of obese mothers has demonstrated that they develop a host of metabolic and neural dysfunctions, with additional evidence of a potential neural mechanism that may be driving the development of these pathophysiologies (Sullivan *et al*., 2012). The current study has demonstrated that HFD exposure during the perinatal period alters the development of hypothalamic-brainstem neurocircuitry, endowing rats with a “stressed” brain-gut phenotype that drives gastric dysfunction, including lower baseline motility, delayed gastric emptying, and an inability to mount a gastric stress response in response to acute stress. Chronically delayed gastric emptying is associated with nausea, bloating, abdominal pain, and early satiety, all of which are exacerbated by dysregulated brain-gut reflexes (Konturek *et al*., 2011). Gastric emptying rates are also an additional, and key, modulator of hunger and satiety, with further effects on meal size and nutrient absorption (Janssen *et al*., 2011). Importantly, the current study has shown that diet alone (in the absence of either offspring or maternal obesity) is sufficient to drive these changes and therefore applicable to all mothers. It remains to be determined whether these developmental alterations, and dysregulated gastric responses to stress, are permanent or whether it is possible they can be reversed with appropriate intervention.

## Data Availability

All data will be made available by the corresponding author upon request.

## Author Contributions

K.E.C., and K.N.B. conceived and designed research; K.E.C., J.A., J.Q.M., and K.N.B. performed experiments; K.E.C., A.T., J.A., J.Q.M., and K.N.B. analyzed data; K.E.C., A.T., and K.N.B. interpreted results of experiments; K.E.C., and K.N.B. prepared figures; K.E.C., A.T., J.A., J.Q.M., and K.N.B. drafted manuscript; K.E.C., A.T., J.A., J.Q.M., and K.N.B. edited, revised manuscript; and approved final version of manuscript.

## Acknowledgements

Supported by NIH grant DK 111667 (KNB) Graphical abstract was created with BioRender.com. We would like to thank F. Holly Coleman for her technical support, particularly with the labeling of gastric-projecting neurons. We would also like to thank W. Nairn Browning as well as Cesare M. and Zoraide Travagli for support and encouragement.

